# Fully integrated SNV, fusion, and isoform analysis of cancer drivers with FLAIR3 for improved functional interpretation

**DOI:** 10.1101/2022.08.01.502364

**Authors:** Colette Felton, Andrea Galvez, Tanvi Damle, Kevin M. Levine, Mark Diekhans, Eunice Lopez Fuentes, Taylor S. Won, Christopher Vollmers, Alejandro Sweet-Cordero, Alice H. Berger, Angela N. Brooks

## Abstract

**Background:** Although the impact of genomic variants and changes in transcriptome regulation are often studied in isolation, their interaction is critical to understanding oncogenesis. We developed FLAIR3, which performs a comprehensive and integrated analysis of single nucleotide variants (SNVs), indels, gene fusions, and alternative splicing using long-read RNA sequencing (lrRNA-seq) and predicts their combined impact on amino acid sequence.

**Results:** By applying FLAIR3 to lrRNA-seq datasets from osteosarcoma and lung adenocarcinoma, we demonstrated that long-read technology captures the full-length molecules required to phase variants with specific isoforms. In the osteosarcoma samples, FLAIR3 identified alternative splicing of gene fusions in known driver genes, including *TP53*. In the lung adenocarcinoma samples, FLAIR3 revealed isoform-biased expression of oncogenic SNVs, including *BRAF*, *CDKN2A*, and *TP53*.

**Conclusions:** FLAIR3 provides a robust method for resolving the complex interplay between variants and transcript isoforms, offering deeper insights into the molecular drivers of cancer.

## Background

Cancer is a disease driven by genomic changes in cells that lead to uncontrollable proliferation. Understanding both the genomic changes and how they cause functional changes to the cells is key to predicting disease prognosis and finding the correct treatment. Next-generation sequencing is already being used in clinical management of cancer and has been shown to improve both progression-free survival and overall survival[1,2]. Presently, the three most common methods of sequencing patient tumors are whole-exome sequencing (allows identification of single-nucleotide variants (SNVs) and small indels in exonic regions)[3], whole genome sequencing (allows identification of SNVs, indels, structural variants (SVs), and copy number changes)[4], panel DNA-based testing (allows identification of SNVs and copy number changes in coding sequence of 100-500 genes), and short-read RNA sequencing (can identify gene expression and some exonic SNVs)[5]. Long-read sequencing allows for the detection of gene expression, alternative splicing, exonic SNVs, and gene fusions[6–9]. However, until recently, long-read sequencing has suffered from lower accuracy than short-read Illumina-based sequencing. With newer methods like PacBio HiFi sequencing, Oxford Nanopore Technologies (ONT) R10 sequencing, and R2C2 + ONT sequencing[10,11], accuracy comparable to or better than Illumina can be obtained[10,12,13]. This allows variants to be detected from long-read sequencing with high accuracy[9,14,15].

SNV detection across large cohorts has revolutionized cancer treatment in tandem with the development of targeted therapies active against the protein products of these recurrent point mutations. However, not all tumors are driven by these common recurrent variants. Novel or rare SNVs can also be detected by short-read approaches, but are often labeled as variants of uncertain significance (VUS), partially due to the difficulty of identifying their impact on protein function.

There are other classes of variants that are more difficult to detect with short-read sequencing, including medium-sized (20-50bp) insertions and deletions (indels) and structural variants that cause gene fusions. Short-read sequencing often fails to detect medium-sized indels due to failed alignment around the indel locus[16]. Short-read sequencing can detect many gene fusions, but tools frequently report high numbers of false positives, and the fraction of shared fusion calls between tools is low[17]. On the genomic level, it has been established that long-read sequencing can detect structural variants much more accurately than short-reads[18,19]. On the RNA level, long-reads have also been shown to have more accurate detection of gene fusions than short-reads[20].

Previous efforts to identify gene fusions in lrRNA-seq data have focused on fusions between two genic regions. However, fusions between a coding gene and a non-coding region can potentially impact the function of the gene as much as a fusion partner with another coding gene. Many cancer fusions involve non-coding sequence[21,22], and detecting these additional fusions could resolve additional drivers.

Some structural variants are uniquely resolvable with long-read sequencing, including complex dense rearrangements where more than two loci are expressed under the same promoter. These complex loci have been previously reported [23,24], but have not been systematically identified. In tumor types prone to complex structural rearrangements such as osteosarcoma, identifying complex fusion loci could also resolve additional drivers.

As with rare SNVs, rare fusions are also often labeled as VUS due to the lack of clinical evidence for their impact. Better predictions of the impact of the variant on protein expression could lead to better ability to characterize and treat rare fusions. Long-read RNA-seq is uniquely able to resolve the full-length molecules expressing the gene fusion, predict the transcript model, and predict the impact of the gene fusion on the expressed amino acid sequence. Previous work has shown alternative splicing of gene fusions [20,24,25], but there has not yet been fusion detection integrated with functional prediction. Previous findings highlight that alternative splicing of gene fusions can alter their tumorigenicity[17,26–28].

For enhanced identification of gene variant detection with lrRNA-seq, we leverage our previous development of the long-read transcript detection and quantification tool FLAIR. FLAIR is a top-performing tool on multiple benchmarks, with a balance of accurate annotated and novel transcript detection[6]. In FLAIR2, we integrated FLAIR transcript detection with SNV haplotype groups to detect transcript bias of variant expression[29]. The ability of FLAIR2 to integrate genotype groups and predict the coding sequence of novel isoforms has been used to identify haplotype association with full-length transcript structure and recategorize SNV interpretation based on transcript assignment[29,30]. Other work in this field has also shown linkage between SNVs and alternative splicing in the same genes[31,32]. However, this analysis is restricted to just SNVs and has not yet been applied to oncogenic alterations. In this work, we developed FLAIR3, which integrates SNV, indel, and fusion detection with transcript detection and quantification and reports protein-coding changes.

Previous work using long-read RNA-seq isoforms to predict protein structure and novel peptides has shown that accurate detection of full-length RNA alterations can reveal important protein domain and localization alterations[33,34], as well as proteomically validated neoantigens[35].

We apply FLAIR3 to sequencing of both osteosarcoma and lung adenocarcinoma to show how integrating variant and transcript analysis enriches the understanding of drivers in both fusion and SNV-driven tumor types. With more complete information about the expressed variant isoforms of cancer driver genes, we can better prioritize variants for treatment with existing therapies or development of novel therapies.

## Results

### FLAIR3 accurately identifies transcript models incorporating fusions and SNVs from lrRNA-seq

Here, we present the FLAIR3 method to report comprehensive variant-aware transcript models from long-read RNA-seq data. FLAIR was originally developed for transcript detection from long-read RNA, and that remains the first step. Previously, FLAIR2 required a specific internal genome alignment (FLAIR align), which produced an additional bed file of reads. This was then passed through FLAIR correct, which produced another bed file of all corrected reads and was then passed through FLAIR collapse to produce a much smaller bed file of isoforms (**Supplementary Fig. 1**). Now, FLAIR3 starts from any aligned bam file, then does correction and collapsing in one step (FLAIR transcriptome), without any bulky intermediate files (**Supplementary Fig. 1**). We also introduce FLAIR combine, which allows users to combine transcriptomes generated from different samples. This provides an alternative analysis path that can be more efficient for large datasets[36].

For fusion isoform detection (**Fig. 1a**), FLAIR fusion extracts chimeric reads from the genome alignment and re-aligns all reads to the reference transcriptome, extracting any additional chimeric reads from that alignment. For fusion detection, FLAIR *fusion* groups chimeric alignments by loci, filtering to those with sufficient read support. It then generates a synthetic fusion reference by combining genomic sequence from all loci involved in the chimeric alignment (**Supplementary Fig. 2**). FLAIR *fusion* is agnostic to the number of genes/loci involved in the fusion, allowing it to detect complex fusions. FLAIR *fusion* then realigns the chimeric reads to the synthetic fusion genome and runs *FLAIR transcriptome* on the synthetic fusion genome (**Supplementary Fig. 3**). In this context, FLAIR3 is not only identifying fusion isoforms, it is also filtering out fusions with inconsistent breakpoints or other artifactual features and stringently identifying the fusion breakpoint location at base-pair resolution. FLAIR *fusion* then takes the preliminary fusion isoform set and further filters the fusions to remove paralogs and promiscuous loci.

**Figure 1:**
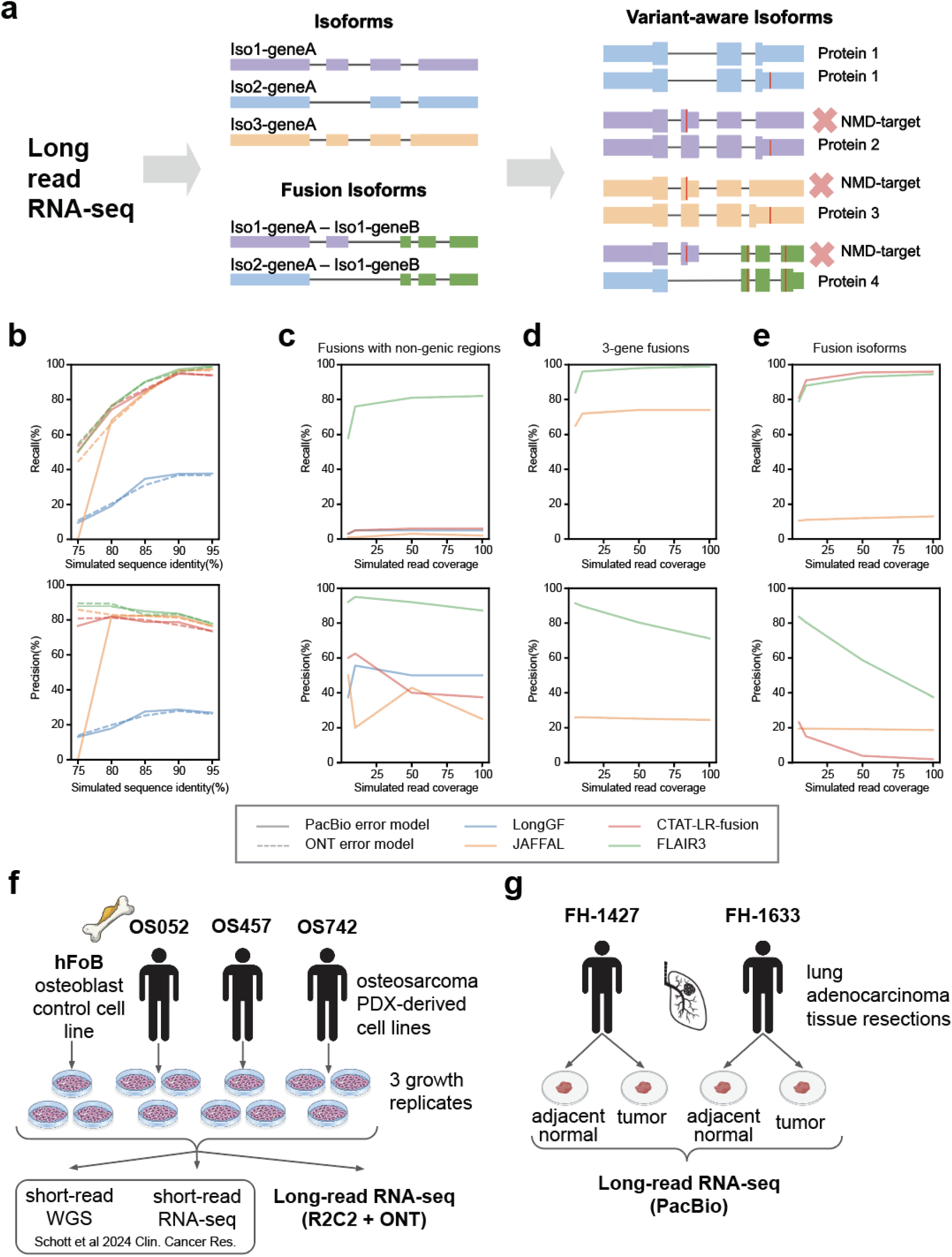
FLAIR3 integrates SNVs and fusions with isoform detection from lrRNA-seq. **a,** FLAIR pipeline: Isoforms, fusion isoforms, and variants are called on each sample individually, then all transcriptomes are combined. Samples are quantified against the combined transcriptome and variants integrated with transcripts. **b,** Recall (top) and precision (bottom) of fusion breakpoint identification (accuracy of within 1bp) on a simulated fusion benchmark with included background data from NA12878 sequencing. **c,** Recall (top) and precision (bottom) of fusion breakpoint identification (accuracy of within 1bp) on simulated fusions with a 5’ gene and 3’ non-genic region. **d,** Recall (top) and precision (bottom) of fusion breakpoint identification (identification of correct genes, JAFFAL only reports genes for 3-gene fusions) on simulated fusions with 3 distinct genes. **e,** Recall (top) and precision (bottom) of fusion isoform identification (correct splice junctions) on simulated fusions. **f,** Cell lines were derived from 3 patient-derived xenograft (PDX) models. Along with a control osteoblast cell line, hFoB. Three biological replicates of each were grown. Each cell line was sequenced by lrRNA-seq using the R2C2 + ONT method. **g,** Tumor resections from two separate patients were split into tumor and adjacent normal tissue then sequenced with PacBio lrRNA-seq. These samples were also clinically genotyped for common cancer variants.

Once FLAIR *fusion* has identified a confident set of fusion transcripts and normal transcripts, they are combined to create the final transcriptome using a new *FLAIR combine* module. Transcriptomes from multiple samples can also be combined with *FLAIR combine* (**Supplementary Figures 1, 3, 4**), which creates a set of unique transcripts identified across all samples. Once the FLAIR transcriptome (FLAIRome) is obtained, each sample can be realigned to the FLAIRome using *FLAIR quantify*, which produces read-to-transcript assignments. Next, the *FLAIR variants* module identifies variant-aware transcripts by matching reads to both variants and transcripts (**Fig. 1a**, **Supplementary Fig. 3-4**)[37]. FLAIR3 takes in a vcf file from a variant caller to do this, and does not perform its own variant calling.

For functional interpretation of variant-aware transcripts, FLAIR3 identifies the coding sequence for each transcript (Methods). It produces tables with quantifications of transcripts, variant-aware transcripts, and predicted amino acid sequences, facilitating differential isoform expression or isoform usage at the transcript isoform, allele-specific isoform, or proteoform level (**Supplementary Figure 3**). This approach facilitates an amino-acid-level functional analysis of differential isoform expression by combining read counts from multiple unique transcript isoforms that would give rise to the same predicted amino acid sequence.

FLAIR transcript identification has been previously evaluated as a top-performing tool for annotated and novel transcripts and was found to identify transcripts with good support from orthogonal data[6]. The FLAIR2-based variant-transcript matching method has also been previously used in multiple studies[29,30]; however, it relied on the variant caller to assign variants to reads. The FLAIR3 *variants* update allows this method to be more agnostic of the variant caller used, so we evaluated DeepVariant[14], longshot[9], Clair3[38], Clair3-RNA[39], and longcallR[40] on PacBio sequencing of human cancer cell lines with matched genomic variant calling from ENCODE (**Supplementary Fig. 5a-b**)[41,42]. We found that DeepVariant, longshot, and longcallR all had consistently high precision and recall, with longcallR having slightly higher precision than any other tool, leading to the highest F1 score. However, all tools except longshot require pre-trained models of the error profile of the sequencing method, as well as informative base-level quality scores. For part of this study, we used R2C2-ONT long-read RNA sequencing, which only longshot was able to successfully call variants on. For this reason, we used longshot in this study, but DeepVariant, longshot, and longcallR are all good candidates for use in generating variant-aware isoforms.

To evaluate fusion detection with FLAIR3, we compared its performance on multiple benchmarks against LongGF[43], JAFFAL[24], and CTAT-LR-fusion[25]. FLAIR3 achieves high precision and recall for fusion breakpoint detection on an established benchmark (**Fig. 1b**)[17]. FLAIR3 slightly outperforms the other long-read RNA fusion detection tools JAFFAL and CTAT-LR-fusion on this benchmark, but overall they all perform comparably. However, FLAIR3 substantially outperforms other tools on recall and precision when it comes to less-standard fusion events, such as fusions with non-genic regions (**Fig. 1c**) and multi-locus fusions (**Fig. 1d, Supplementary Fig. 5c**). Multi-locus fusions are not identified by CTAT-LR-fusion[25] and while JAFFAL[24] reports putative 3-locus fusions, it does not report breakpoints for them.

FLAIR3 also outperforms other fusion detection tools for the identification of fusion isoforms. JAFFAL only reports one isoform per fusion breakpoint, so its recall is low, while CTAT-LR-fusion has high recall but little filtering of fusion isoforms, resulting in low precision. FLAIR3 balances recall and precision to achieve high accuracy in detecting fusion isoforms, including both fusions with splicing away from the breakpoint (**Fig. 1e**) and fusions with alternative splicing causing multiple apparent fusion breakpoints (**Supplementary Fig. 3c**).

### Application of FLAIR3 to osteosarcoma and lung adenocarcinoma lrRNA-seq data

We applied FLAIR3 to two sample sets with different sequencing methods and tumor mutational profiles (**Fig. 1f**). The first set used three osteosarcoma patient-derived xenograft (PDX)-derived cell lines that were previously characterized using short-read WGS and short-read RNA-seq[44,45]. From WGS analysis, putative driver alterations were identified for each cell line. With this set, we are able to compare variant analysis derived from long-read RNA-seq to more traditional methods of variant detection. Additionally, osteosarcomas are known to be driven by complex structural rearrangements like chromothripsis, which we hypothesize would produce more complex fusions at the RNA-level that are undetectable with short-read methods[46]. We performed long-read RNA-seq using R2C2 + ONT (**Fig. 1f**). We obtained a median read length of 1732bp and 99.4% sequence accuracy with this sequencing method (**Supplementary Fig. 6a**).

To demonstrate that the utility of FLAIR3 is agnostic to the sequencing platform, we also performed PacBio Kinnex Iso-Seq on two primary human lung adenocarcinomas and matched tumor-adjacent normal tissue samples (**Fig. 1g**). With this sequencing method, we obtained a median read length of 1533bp and 99.7% sequence accuracy (**Supplementary Fig. 6b**).

### Long-read RNA-seq can identify gene alterations identified by WGS in osteosarcoma

The osteosarcoma PDX-derived cell lines were previously characterized through WGS to identify copy number changes, structural variants, SNVs, and indels in clinically actionable genes[44,45]. We ran FLAIR3 on these samples to identify amino acid sequence usage differences in each PDX-derived cell line compared to a more normal osteoblast control cell line (hFoB), whether due to structural variants, SNVs, splicing, or a combination. For comparison with other lrRNA-seq fusion detection methods, we also called fusions with LongGF, JAFFAL, and CTAT-LR-fusion. We also identified changes in gene expression relative to the normal control. Consistent with osteosarcomas primarily being driven by structural rearrangement, we identified both gene expression changes associated with copy number amplification and gene fusions driven by structural variation (**Supplementary Fig. 7a**).

Using FLAIR3 on matched lrRNA-seq, we were able to identify associated gene expression changes for genes with copy number changes in WGS. *CCNE1*, *MYC*, and *CDK4* were identified as amplified drivers from WGS[44,45] and we observed significant overexpression of these genes in the corresponding PDX-derived cell lines. *PTEN* was identified as a driver due to copy number loss in OS052. We observed a loss of expression in OS052, but due to low expression in the control osteoblasts, it was not identified as a significant event from the lrRNA-seq.

In OS457, which is driven by the *CCNE1* amplification, FLAIR3 identified a fusion between the 3’ UTR of *CCNE1* and a pericentromeric region of chr19. This demonstrates that lrRNA-seq analysis can provide more detailed characterization of the transcriptomic consequence of a genomic alteration (**Fig. 2a-b**). FLAIR3 was the only lrRNA-seq fusion caller to identify this *CCNE1* fusion and the fusion was also missed by the published WGS analysis. Visual inspection of read support in WGS and short-read RNA-seq (RNA-seq) validates the FLAIR3 findings (**Fig. 2c**).

**Figure 2:**
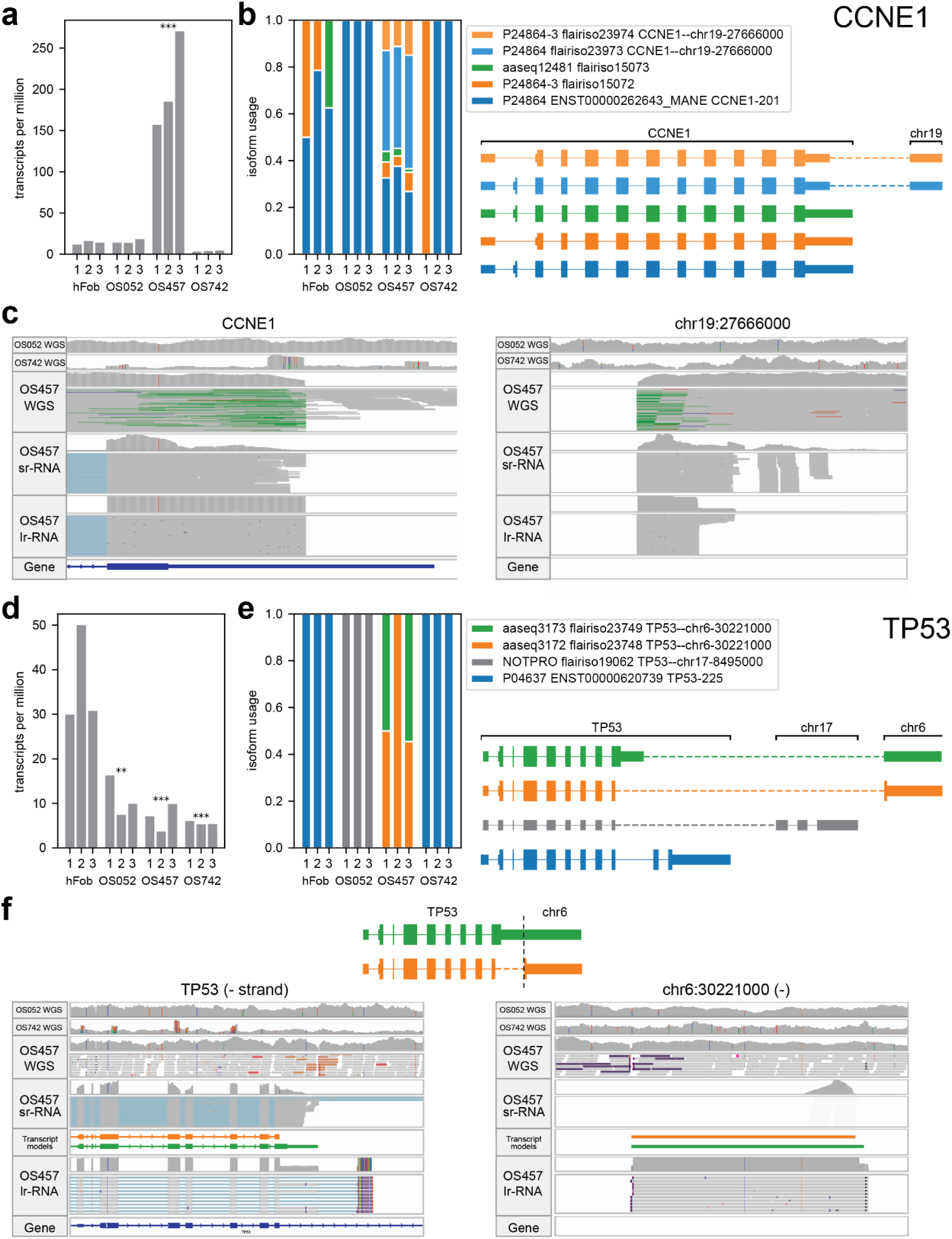
FLAIR3 detects additional alterations in osteosarcoma drivers detected with WGS. **a,** Gene expression (transcripts per million, TPM) for *CCNE1*. Significance relative to hFoB (*=p<0.05, **=p<0.01, ***p<0.001). **b,** Isoform usage for *CCNE1*. x-axis is cell line, then biological replicate. Left: Isoform usage (fraction of total expression of the gene). Different shades of the same color represent isoforms that produce the same amino acid sequence. Grey means they are unproductive. Legend: Each isoform is identified by the UniProtKB accession (when possible), then Ensembl transcript ID and transcript name (when possible). MANE transcripts are indicated. If the isoform is of a gene fusion, the gene ID is in the form geneA--geneB. Right: Models of the isoforms. All are oriented in the 5’->3’ gene direction.Thicker blocks are coding sequence (CDS). **c,** Raw read alignments at the fusion breakpoint for the OS457 *CCNE1*–chr19:27666000 fusion. Top: WGS, middle: short-read RNA-seq, bottom: long-read RNA-seq (fusion isoform models included in black). Colored reads indicate non-matching pair orientation. Red bar at end of alignment indicates additional sequence not aligned to this locus. **d,** Gene expression for *TP53*. Significance is as described in a. **e,** Isoform usage for *TP53*. All panels are as described in b. **f,** Inferred model of *TP53* fusion transcript structure using both the RNA and WGS data and raw read alignments at the fusion breakpoint for the OS457 *TP53*–chr6:30221000 fusion. Tracks are as described in c.

We found that over 55% of *CCNE1* expression is driven by this fusion (**Fig. 2b**). Given that from WGS, *CCNE1* was found to be expanded to 42 copies in OS457[44], this indicates that either this specific fusion locus is much more highly expressed or that one haplotype of *CCNE1* forms this fusion and is then amplified along with the pericentromeric region. Consistent with the latter, WGS read coverage near the fusion breakpoint in both the *CCNE1* locus and chr19 locus suggests that this fusion represents the amplified copy of *CCNE1* (**Fig. 2c**). We also found that the *CCNE1* fusion transcript isoforms have a similar splicing pattern to the non-fusion CCNE1 transcripts, in which they include or skip the second exon, suggesting that this fusion is not impacting the alternative splicing of *CCNE1* (**Fig. 2b**). Although there is no significant change in *CCNE1* splicing, we do note that flairiso15073, which has an alternative acceptor in exon 4 and produces a distinct protein, is expressed only in the non-fusion reads.

We also observed *TP53* fusions in OS052 and OS457 (**Fig. 2d-e**). These fusions were both reported from short-read WGS analysis[44,45](**Fig. 2f**). The OS052 *TP53* fusion was found by LongGF, but missed by JAFFAL and CTAT-LR-fusion. The OS457 fusion was not reported by any other long-read RNA fusion caller. This lack of detection by other RNA-based tools may be due to both fusion breakpoints occurring in non-exonic regions (OS052 breakpoint is intronic, OS457 breakpoint is intergenic).

In addition to being able to recover these fusions, which are missed by many other approaches, we identified both the splicing and functional consequences of these fusions (**Fig. 2e**). The OS052 *TP32*–chr17 fusion introduces an early stop codon and downstream splice junctions (>50nt), so is predicted to undergo NMD and not produce protein. In contrast, the OS457 *TP53*–chr6 fusion also introduces an early stop codon, but is not spliced beyond that stop codon, indicating that it likely produces a truncated protein product. Additionally, the *TP53*–chr6 fusion has two isoforms that contain distinct apparent fusion breakpoints (**Fig. 2f**). When comparing the apparent fusion breakpoints with the genomic breakpoint from the WGS data, we can see that one isoform (flairiso23748) splices out the sequence between the *TP53* splice site and the 3’ fusion breakpoint on chr6, while the other retains that sequence. This shows how lrRNA-seq analysis is able to enhance gene fusion characterization and provide more detailed functional information than WGS alone.

### FLAIR3 can identify novel alterations in osteosarcoma

FLAIR3 also identifies novel alterations that are not identified from WGS. FLAIR3 identifies additional fusions in *NOTCH1*, *NOTCH3*, and *SMAD2*, but only the *NOTCH1* and *NOTCH3* fusions are predicted to produce protein products. The fusion between *NOTCH3* and a nongenic region on chr19 is predicted to produce a protein that contains solely EGF-like repeats that form the extracellular portion of NOTCH3, with none of the sequence of the fusion partner expressed (**Fig. 3a-b**). Although this fusion does not express the intracellular domain of NOTCH3, which functions as a transcription factor, previous work has shown that the extracellular domain of NOTCH proteins can regulate the NOTCH pathway[47].

**Figure 3:**
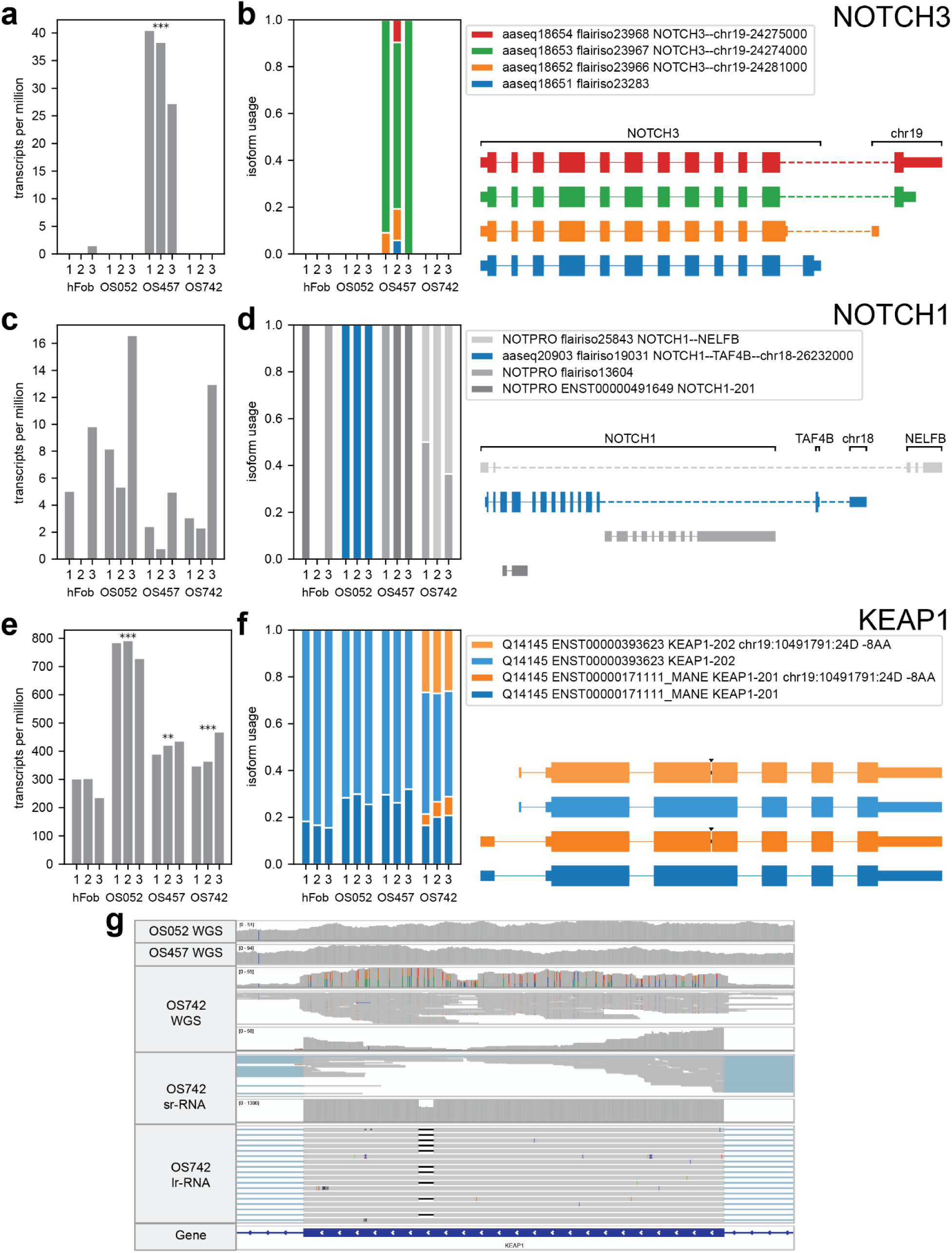
FLAIR3 detects novel variants in cancer genes in osteosarcoma that cannot be detected with short-read methods. **a,** Gene expression for *NOTCH3*. Significance is as described in Figure 3a. **b,** Isoform usage for *NOTCH3*. All panels are as described in Figure 3b. **c,** Gene expression for *NOTCH1*. Significance is as described in Figure 3a. **d,** Isoform usage for *NOTCH1*. All panels are as described in Figure 3b. **e,** Gene expression for *KEAP1*. Significance is as described in Figure 3a. **f,** Isoform usage for *KEAP1*. All panels are as described in Figure 3b. **g,** Raw read alignments around the deletion in *KEAP1* in OS742. Top: WGS, middle: short-read RNA-seq, bottom: long-read RNA-seq.

We also identified a three-locus fusion between *NOTCH1*, a *TAF4B* exon, and a *TAF4B* intron on the opposite strand (chr18-26232000, **Fig. 3c-d**). Similar to the *NOTCH3* fusion, this fusion is predicted to produce a protein that contains the extracellular repeats of NOTCH1, along with a small amount of additional sequence from TAF4B. LongGF, JAFFAL, and CTAT-LR-fusion also detect a *NOTCH1*--*TAF4B* fusion, but only FLAIR3 reports the full complexity of the locus. Additionally, complex loci like this can only be fully resolved using long-read sequencing and have likely been missed in previous studies that use short-read WGS.

We also report a novel deletion in *KEAP1* in OS742. KEAP1 represses NRF2 and its dysregulation can lead to oncogenesis and greater resistance to therapies[48]. This alteration is a 24bp deletion that causes an in-frame 8 amino acid deletion (KEAP1 p.G371_V378del, deletes GGLLYAVG, **Fig. 3e-f**). This deletion is undetectable through either short-read WGS or short-read RNA-seq, as the reads fail to align around the deletion area (**Fig. 3g**). This is likely due to incorrect alignment of the WGS reads near the deletion, which is a known issue for deletions in the 20-50bp range[16]. Using the predicted amino acid sequence from FLAIR3, we were able to model the KEAP1 protein structure and find that this deletion causes a slight change in the protein, although the functional consequence is not known (**Supplemental Fig. 8a**). Since KEAP1 represses NRF2 protein activity, we investigated the expression of genes downstream from NRF2 (**Supplemental Fig. 8b**)[49]. We found no differential expression of NRF2 target genes compared to the osteoblast control; therefore, additional work is needed to determine if the deletion is functional or a passenger mutation. Overall, these data show our ability to identify multiple types of additional cancer-relevant variants in these osteosarcoma samples, as well as being able to add additional splicing context to known alterations.

### Identification of tumor-biased isoform changes in lung adenocarcinomas

While osteosarcomas are driven by structural rearrangements, lung adenocarcinomas have one of the highest SNV mutation rates in cancer[50] and are frequently driven by mutations in key RAS pathway genes. 84% of lung adenocarcinomas have RAS activation and over 50% are driven by mutations in *KRAS*, *BRAF*, and *EGFR[51]*. Other recurrent drivers in tumors without a mutation in those 3 genes are *KEAP1* and *TP53[52]*. In our lung dataset, we have matched tumor-normal samples, which enable investigation of tumor-specific splicing changes.

FLAIR3 analysis of primary tumors against matched normals identified predicted coding sequence changes due to both variants and splicing. In clinically relevant druggable genes (n=150), we identified three damaging somatic SNVs and 21 splicing changes in FH-1427 and 1 damaging somatic SNV and 29 splicing changes in FH-1633 (**Supplementary Fig. 7**). In these samples, we did not identify any fusions in actionable cancer genes with >10% usage. We noted that while the SNVs were unique to each patient, many of the splicing changes observed were shared between the tumor samples. This includes splicing in *ERG*, *SYK*, *XPO1*, *MITF*, *MDM2*, *SMARCB1*, *RAD21*, *ERCC1*, and *DNMT3A* (**Supplementary Fig. 7, Supplementary Fig. 9**).

Since FLAIR3 is reporting at the level of amino acid sequence changes, all of those splicing changes are predicted to change proteoforms in the cell. For example, we observe that *ERG*, an oncogenic transcription factor, had decreased expression coupled with decreased usage of the MANE isoform (ERG-201) in both lung samples (**Supplementary Fig. 9a-b**). Both samples show increased usage of isoforms using upstream transcription start sites (TSSs) (ERG-206, ERG-207, ERG-208, and ERG-209), leading to increased usage of the P11308-1 and P113080-3 proteins, which contain different amino acids at the N-terminus when compared to the canonical protein. Although alternative splicing in *ERG* has been shown to alter oncogenicity, this TSS shift has not been previously characterized[53]. Although a larger cohort would be needed to confirm these splicing changes, FLAIR3 accelerates identification of changes at the protein level from long-read RNA.

### Identification of alternative isoform expression and allele bias of driver variants

In sample FH-1633, the only damaging somatic variant we identified was the canonical driver variant BRAF V600E, which had an interesting association with the isoform expression of BRAF (**Fig. 4a-b**). The genomic position of the BRAF V600E variant is present on both the BRAF-220 (MANE) and the BRAF-204 isoforms. However, we only see the variant expressed on the BRAF-220 isoform, indicating isoform-specific allele expression (Fisher’s exact test, p=0.004). We also observe that total usage of the BRAF-220 isoform almost doubles from 36% in the normal sample to 64% in the tumor sample and all of that increase in usage can be attributed to reads containing the V600E variant. Identifying that the V600E variant is expressed exclusively on the MANE isoform of BRAF allows us to conclude that it is likely performing its canonical role in this tumor sample.

In sample FH-1427, although we did not detect any canonical RAS driver variants, we identified damaging somatic variants in *KEAP1*, *TP53*, and *CDKN2A*. Although none of these events is a canonical RAS driver, these are all genes that have been previously identified as recurrently mutated in lung adenocarcinomas[54]. 1.3% of lung adenocarcinoma patients in the GENIE study had a variant in all 3 genes[55,56]. Based on the data in GENIE, *CDKN2A* variants significantly co-occur with both *TP53* and *KEAP1* mutation (both p<0.001, GENIE v18.0)[55].

*TP53* expresses two primary isoforms in FH-1427: TP53-225 and flairiso37824 (**Fig. 6c-e**). TP53-225 produces the canonical p53 protein (p53, P04637), while flairiso37824 has a downstream TSS, uses a downstream start codon, and produces a truncated version of p53 (Del133-p53, P04637-7). Del133-p53 lacks the N-terminal p53 TAD domains, while retaining most of the DNA binding domain and dimerization ability. It has previously been shown to repress transcriptional impacts of p53 and be associated with cancer[57].

**Figure 6:**
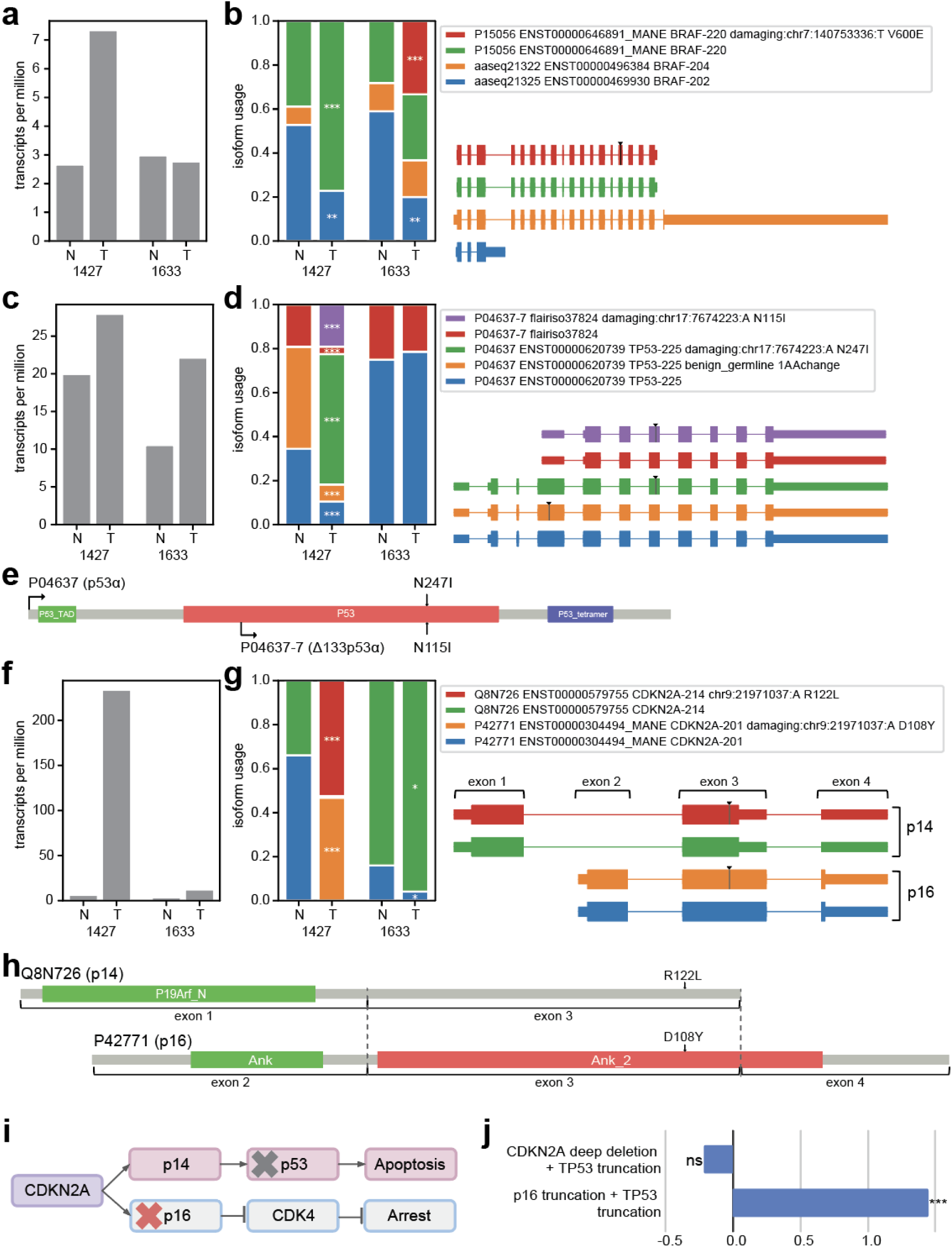
Damaging somatic variants have isoform-specific expression and isoform-specific impacts. **a,** Gene expression for *BRAF*. **b,** Isoform usage for *BRAF*. All panels are as described in Figure 5b. **c,** Gene expression for *TP53*. **d,** Isoform usage for *TP53*. All panels are as described in Figure 5b. **e,** TP53 protein structure with the somatic mutation from **d** annotated. The protein starts for the isoforms in **d** are also annotated. **f,** Gene expression for *CDKN2A*. **g,** Isoform usage for *CDKN2A*. **h,** CDKN2A protein structure for the p14 and p16 isoforms (out of frame with each other), with dashed lines showing where they share genomic sequence. The somatic mutation from d is annotated. **i,** Simplified diagram of the CDKN2A pathway, showing the distinct function of the p14 and p16 protein isoforms. Red X shows proposed impact of variant from panel h. Grey X shows proposed impact of variant from panel d. **j**, Bar plot of log2 odds ratio of co-occurrence in GENIE LUAD of different CDKN2A mutations with TP53 truncation.

In the tumor-adjacent normal tissue, we observe that FH-1427 has a predicted benign germline variant in TP53. Given that this variant is only in the CDS of the TP53-225 isoform, we only report it on TP53-225, with its expression making up about 50% of that isoform (**Fig. 6c-d**). In the tumor sample, we detect a damaging somatic variant. This variant is expressed on both isoforms; however, we see a biased allele expression of the somatic variant with the canonical p53 protein (TP53-225 usage vs flairiso37824).

We also detect a damaging somatic variant in *CDKN2A*. This gene also has strong upregulation in the tumor sample (**Fig. 6f**). In the FH-1427 tumor-adjacent normal tissue, we see two isoforms expressing only reference sequence: CDKN2A-201 (MANE, 66% usage) and CDKN2A-214 (34% usage). In the tumor sample, we observe allele-biased expression in *CDKN2A* in which all transcripts express the damaging somatic variant (**Fig. 6g**). This full penetrance of variant expression suggests that the haplotype of *CDKN2A* with the variant is being amplified or stabilized. Additionally, the other haplotype may be silenced or deleted. We also observe a switch in the dominant isoform in the tumor sample, with CDKN2A-214 increasing to 52% usage.

CDKN2A-201 and CDKN2A-214 have different first exons and different annotated start codons. Their reported start codons are out of phase and they express two distinct amino acid sequences. CDKN2A-201 produces the canonical p16 (P42771) protein, which regulates the cell cycle through repressing CCND1 and CDK4[58]. CDKN2A-214 produces the p14 (ARF, Q8N726) protein, which activates p53 through repressing MDM2[59].

Since CDKN2A-201 and CDKN2A-214 have distinct ORFs, the damaging variant impacts different portions of each protein (**Fig. 6h**). On CDKN2A-201, the SNV (D108Y) impacts the ankyrin repeat domain which binds CDK4. However, on CDKN2A-214, the SNV (R122L) is near the C-terminal of the protein and not in a functional domain. Therefore, this variant may have more impact on the function of p16 than p14 and may reduce the ability of *CDKN2A* to regulate the cell cycle through CDK4, while maintaining its ability to regulate TP53 (**Fig. 6i**).

The coupling of this variant that may lead to only the p14 protein being functional with increased usage of the p14-producing isoform may indicate that this variant shifts cells into a state where they are more dependent on p53 to regulate the cell cycle. Since FH-1427 also has a damaging variant in *TP53*, the knockdown of *TP53* and *CDKN2A* together may work to dysregulate the cell cycle and promote tumor growth.

We investigated further the interaction between *TP53* and *CDKN2A* mutations using the GENIE v18.0 LUAD cohort. Although the mutation we observed in FH-1427 was missense, we selected for truncations in the GENIE cohort to be confident that the mutations knocked out p16 function. We found that SNVs that truncate the p16 protein isoform of *CDKN2A* co-occur significantly with *TP53* truncation (chi^2^ p-value <<0.001, log2 odds ratio 1.45, **Fig. 6j**). In contrast, *TP53* truncations do not significantly co-occur with homozygous deletions of *CDKN2A* that knock out the function of p16 and p14 (chi^2^ p-value=ns, log2 odds ratio -0.21). This further supports that the CDKN2A p14 and p16 isoforms have distinct tumor suppressive roles and that SNVs should be considered in terms of their impacts on both isoforms to interpret their functional impact. We show that an integrated analysis of gene fusions, SNVs, and indels with RNA processing not only allows recovery of more actionable cancer variants, it also enriches our understanding of the functional impact of the variants identified.

## Discussion

In this study, we developed and benchmarked FLAIR3 to use lrRNA-seq to accurately detect fusion isoforms and integrate SNVs with splicing in clinically actionable cancer genes. We additionally predict the functional impact of combined genomic and transcriptomic variation in PDX-derived cell lines and primary tumor samples. Like previous versions of FLAIR, the FLAIR3 analysis pipeline is made publicly accessible to other researchers. The integrated analysis of SNVs, indels, fusions, and isoforms on the same reads not only allows us to effectively investigate more cancer types, it also allows an unprecedented ability to link these alterations together for a more functional analysis of the impact of the mutations on the tumor. This allows for better prioritization of variants by understanding the level at which they knock out or enhance gene function. For instance, the variant we observed in *CDKN2A* is only predicted to alter the function for one isoform of the gene.

With FLAIR3, we automate fusion isoform detection and functional prediction, thus improving fusion detection over existing methods. Alternative splicing of gene fusions has been underexplored, yet they could be important sources of tumor-specific neoantigens. For example, we identified fusion isoforms in *TP53* and *NOTCH3* that are predicted to create novel tumor-specific peptides, which would not be reported by existing fusion detection tools. We also systematically detected and benchmarked fusions across >2 loci, which led to the identification of the *NOTCH1–TAF4B(+)*–TAF4B(-) fusion in osteosarcoma. This is another type of event that is underexplored in its impacts on tumorigenesis. Other difficult-to-resolve and underexplored variants that FLAIR3 detects include medium-sized indels such as the 24bp deletion we identified in *KEAP1*.

Our method of focusing on functional changes at the amino acid sequence level allows us to prioritize isoform changes and rapidly identify those that change important proteins in cancer. Many splicing and RNA processing changes only affect the UTR of a gene or produce a transcript that is likely to be degraded by NMD. There are also many cases where multiple splicing events result in the same change to the amino acid sequence. By performing our analysis on the inferred amino acid sequence, we both prioritize and increase sensitivity to protein-coding changes.

Identifying the isoform context of a variant can also have implications for predicted drug response. For instance, we identified that the *BRAF* V600E variant in sample FH-1633 is expressed on the BRAF-220 (MANE) isoform but not on the BRAF-204 isoform. However, previous studies[60] have shown that the BRAF-204 is actually much more highly expressed than BRAF-220 in many tumor types. Little is known about how response to BRAF inhibitors may change depending on isoform-specific expression of BRAF variants, but this evidence suggests that this gene is capable of producing a diversity of protein products and that their response to BRAF inhibitors should be studied further.

Long-read RNA-seq technology has come a long way in the past few years. Increased confidence in this method is due to the increased read accuracy and read lengths that ONT and PacBio sequencing protocols have recently achieved. However, as with any sequencing method, there are still biases. Long-read cDNA libraries generally have a maximum read length of about 3kb due to the limitations of RT-PCR. This means that this method cannot detect splicing or variants in very long transcripts. This method is also dependent on high-quality libraries with long median read lengths, because we retain only reads that support full-length transcripts for our quantification of variant-transcript interactions. This means that in libraries with a high number of truncated reads, only a fraction will be informative for this analysis.

Also, although this method predicts productivity and amino acid sequence based on the RNA sequence, there are many complex factors that influence which proteins are generated and stabilized in cells. For instance, although NMD has characteristic transcript features, there are also many known transcripts that exhibit NMD-escape[61]. Although this method incorporates known NMD-escape transcripts, we cannot predict novel NMD escape and thus may be slightly under-representing productive transcripts.

## Conclusions

We believe that the FLAIR3 method and its holistic approach to identifying the functional impacts of gene alterations has promise not only for a deeper understanding of the interaction between variants and RNA processing, but also as a clinical tool for improving patient outcomes.

## Methods

### Experimental Design

The objectives of this study were to comprehensively identify many types of cancer driver alterations in multiple cancer types from long-read RNA-seq and reveal interactions between genomic and transcriptomic alterations in cancer drivers. We developed tools for long-read RNA sequencing analysis and validated them using simulated data benchmarks and publicly available cell line sequencing data. To examine a range of driver alterations, we selected further study in osteosarcoma, which is primarily driven by structural variants and lung adenocarcinoma, which has canonical driver alterations due to SNV, fusions, and splicing. We performed computational analysis on long-read R2C2 cDNA + ONT sequencing of 3 biological replicates of osteosarcoma patient-derived cell lines and compared each to sequencing of an osteoblast control to identify significant events. We then compared our results to short-read WGS and short-read RNA-seq from Schott et al. We then further performed computational analysis on PacBio cDNA sequencing of two pairs of matched tumor and normal primary samples from lung adenocarcinoma. Osteosarcoma patient-derived cell lines are described in Schott et al[45] and were obtained from the Sweet-Cordero lab at UCSF. Primary lung samples were obtained from surgical resections performed at Fred Hutchinson Cancer Research Center.

### Growing osteosarcoma cell lines and extracting and evaluating RNA

The cell lines used were hFoB, OS052, OS457, and OS742[44]. They were cultured in DMEM with 1% Pen/Strep and 10% FBS. They were split using Gibco trypsin-EDTA at 0.05% for 7-10 minutes. We used the Zymo Direct-zol RNA Miniprep Kit (#R2050) to isolate RNA from TriZol, then evaluated the RNA quality using a NanoDrop One, 4200 TapeStation System, and Qubit Fluorometer 3000. All samples used had RIN scores above 9. 2ug of RNA were used for sequencing.

### Library preparation and preprocessing osteosarcoma data with the R2C2 method

Libraries were prepared and sequenced using the R2C2 protocol as described in Adams et al[10,11]. Briefly, reads were sequenced with ONT R10 flow cells, basecalled with Guppy 6.5.7 sup model, then consensus called with C3POa v3.1 and demultiplexed with C3POa_postprocessing. PolyA tails were removed with a Mandalorion (v4.3.1) utility and reads were merged based on UMIs. Error rate was calculated by samtools stats (cigar) from alignment to hg38[62]. This resulted in an average read accuracy of 99.7%.

### Lung Cancer Specimens

Tumor and adjacent normal lung tissue specimens were obtained from surgical resections performed as part of standard clinical care. Adjacent normal tissue was defined as tissue grossly free of tumor involvement at the time of resection, as assessed by the operating surgeon and confirmed by visual inspection in the pathology suite. After tumor specimens were sent for standard clinical processing, excess tissue was snap-frozen in liquid nitrogen for purposes of this research study. All samples were collected in accordance with institutional protocols and ethical guidelines, with informed consent obtained from all patients under an Institutional Review Board (IRB)-approved protocol at the University of Washington / Fred Hutchinson Cancer Center (RG1000720).

### Lung sample RNA Extraction

Total RNA was extracted from tissue samples using the AllPrep DNA/RNA Mini Kit (Qiagen, Cat. No. 80204), following the manufacturer’s protocol. Briefly, up to 30mg of frozen tissue was homogenized in Buffer RLT containing β-mercaptoethanol using a mechanical tissue homogenizer. After a series of wash steps, RNA was eluted in RNase-free water and samples stored at –80°C until further use. An additional on-column DNase digestion was performed using the Qiagen RNase free DNase set (Cat. No. 79254).

### Lung sample Kinnex Full-Length Library Preparation

Kinnex full-length sequencing libraries were prepared using the Kinnex full-length RNA Kit (PacBio, Menlo Park CA) according to the manufacturer’s instructions. Briefly, quality of the input RNA was determined by measuring RNA Integrity Number (RIN) with the RNA ScreenTape and Reagents using a 4200 TapeStation System (Agilent, Santa Clara, CA) to confirm that all samples had RIN values ≥7.0, and the quantity of the input RNA was determined using a Lunatic UV/Vis spectrophotometer (Unchained Labs, Pleasanton, CA). A total of 300 ng of total RNA for each sample was then carried forward into cDNA synthesis, where each sample used a uniquely barcoded cDNA primer. cDNA yields were quantified using a Qubit Fluorometer and the 1x dsDNA HS Kit (Thermo Fisher Scientific, Waltham, MA) and cDNA fragment sizes were confirmed with the High Sensitivity D5000 ScreenTape and Reagents using the Agilent 4200 TapeStation System. The barcoded cDNA from each sample were subsequently pooled in equal mass ratios and then split across eight parallel Kinnex PCR reactions to add the adapters necessary for forming the final 8-mer Kinnex array. Equal volumes of these Kinnex PCR products were then pooled and carried through the Kinnex array formation steps using a single barcoded Kinnex adapter. The concentration of the final Kinnex array was determined using a Qubit Fluorometer and the 1x dsDNA HS Kit and the expected size of the Kinnex array was confirmed with the Genomic DNA ScreenTape and Reagents using the Agilent 4200 TapeStation System.

### Long-Read RNA-Sequencing of Lung Cancer Samples

PacBio HiFi data were generated per manufacturer’s recommendations at the University of Washington Long Reads Sequencing Center. Libraries were quality checked for concentration (Qubit dsDNA HS (Thermo Fisher, Q32854) measured on DS-11 FX (Denovix)) and size distribution (FEMTO Pulse (Agilent, M5330AA & FP-1002-0275)). The libraries were sequenced on the PacBio Revio platform on one SMRT Cell 25M with Revio Chemistry V1 (PacBio, 102-817-900) using Adaptive Loading and a 30-hour movie. Data were postprocessed using SMRT Link v13.1 with the “Read Segmentation and Iso-Seq” pipeline to segment, classify, and demultiplex the reads.

### FLAIR3 computational pipeline (isoforms + quantification)

FLAIR v3.0.0b1 was used in this study. GENCODE v38 was used as the transcript reference[63]. All samples were processed as in Supplementary Fig. 1, starting with stranded reads from upstream processing pipelines. For osteosarcoma data, all 3 replicates of each cell line were combined for isoform/fusion/SNV identification. For lung data, isoform/fusion/SNV detection was performed on each sample individually. For both datasets, the transcriptomes (including fusions) were then merged and each dataset was quantified through realignment of individual samples to the merged transcriptome (FLAIR quantify with --check_splice, --stringent, --output_bam).

### Statistical comparison of gene expression changes

Gene expression was calculated from FLAIR-quantify output. We combined counts for all isoforms for a given gene to get the gene expression, then normalized by counts per million. For the osteosarcoma data, we ran DESeq2[64] through FLAIR-diffexp, comparing the replicates of each patient-derived cell line to hFoB to identify significantly altered genes. For the lung cancer data, we calculated the log2 fold change for each gene between the tumor and paired normal sample. We did not calculate p-values for gene expression changes in this data.

### FLAIR fusion computational pipeline

As described in Supplementary Fig. 1, FLAIR fusion takes in all chimeric (with SA tag) reads from genomic alignment with minimap2 2.28[65] (minimap2 -ax splice --secondary=no -s 40 -G 350k GENOME.fa READS.fa). It then further aligns all reads in the sample to the reference transcriptome and identifies chimeric reads from transcriptomic alignment. It then combines the genomic and transcriptomic putative chimeras based on genomic breakpoint location and filters out those supported by only one read and those where the most 5’ alignment position is >15kb from an annotated transcript start. It then takes the remaining chimeras and generates a synthetic fusion reference by taking the outer genomic bounds of the existing alignments to each locus and concatenating the genomic sequence therein based on the putative breakpoint (+1kb). FLAIR fusion also generates annotation for the synthetic genome through converting reference annotations for the locus into synthetic genome coordinates and combining annotations across loci. It then realigns all chimeric reads to the synthetic fusion genome with minimap2 -uf, which does not force canonical splice junctions. Next, FLAIR fusion detects novel splice junctions/fusion breakpoints through identifying positions sufficiently distant from annotated splice junctions with sufficient read support. It then runs FLAIR correct to adjust splice junctions using the synthetic genome-aligned reads, the synthetic annotation, and the novel consensus junctions. Next, it runs FLAIR collapse with –check_splice and –stringent to identify isoforms aligned to the synthetic genome, which enforces good alignment around each splice site and thus enforces consistent fusion breakpoints. The final step, which converts from synthetic aligned transcripts to reference genome-aligned transcripts and generates the final fusion set, first filters for only transcripts that cover all loci involved in the fusion. Additional filters include: removing fusions with genes that are too promiscuous (involved in more than 4 fusions), fusions with only one read, fusions between paralogs, fusions with mitochondrial genes, and fusions with immunoglobulin genes.

### FLAIR variants computational pipeline

For variant-isoform assignments, FLAIR variants takes in a VCF of variant calls along with a filtered bam file of full-length alignments to the transcriptome (from FLAIR quantify) for each sample. It then combines variants from all samples and converts them to transcriptomic coordinates. For each read aligned to the transcriptome, FLAIR variants identifies the set of variants present on that read. For SNVs, it only uses positions referenced in the VCF. For indels, it recognizes any insertion or deletion >10bp in the read. Then for the set of reads aligned to each transcript, it identifies confident variant sets by first: identifying all individual variants with read support >= 3 on that transcript, then filtering variants present on reads to only those with sufficient support across the transcript, then grouping reads based on shared variant sets. FLAIR variants then generates an expanded variant-aware transcriptome where each isoform + variant set is reported separately.

In this study, we focus on applications of FLAIR3 to cancer genomic studies; therefore, we filtered for variants to those that were likely somatic as classified in GENIE[55] and probably damaging as determined by PolyPhen[66]. We then further filtered alterations to those occurring in actionable-druggable cancer genes[44].

### FLAIR productivity and amino acid sequence prediction

From this transcriptome, we can then predict the productivity of isoforms. We identify start codons using only those annotated in GENCODE, then detect stop codons based on transcript sequence. We select the longest detected ORF on each transcript. Transcripts are predicted as either productive (PRO), no start codon (NGO), no stop codon (NST), or early stop codon (NMD, stop codon is earlier than 55bp before the end of the second-to-last exon[67]). Once we have identified the CDS of the FLAIR transcripts, we can predict the amino acid sequence of each transcript. An additional step then takes in the variant-aware isoforms and original CDS predictions and adjusts them based on the sequence alterations caused by the variants. We then report additional files where for each amino acid sequence, we report all isoforms that produce this sequence. For amino acid sequence quantification, we combine the reads that support each isoform that produces this sequence. Amino acid sequences that match UniProt[68] protein sequences are annotated with the corresponding UniProtKB accession number.

### Statistical comparison of amino acid sequence changes

For the osteosarcoma data, we ran DRIMSeq[69] through FLAIR-diffexp on the quantification of the predicted amino acid sequences, comparing the replicates of each patient-derived cell line to hFoB, to identify significantly altered amino acid sequences. For the lung cancer data, we ran the FLAIR script diff_iso_usage on each tumor normal pair. Here significance is determined using a Fishers exact test.

### Actionable-druggable gene list

This is a list of 252 genes defined as clinically actionable and druggable as defined in Sayles et al[44].

### Filtering of results based on druggable genes and damaging variants

We labeled each predicted amino acid sequence based on whether it matches any annotated UniProt sequence as well as whether it contains any variants that overlap with the GENIE project somatic variant calls[55]. We then filtered for sequences that either made up >=5% of gene expression and had >=3 reads or had >=50% of gene expression and >= 2 reads. We then re-quantified isoform usage based on only those sequences. To identify significant events, we only looked at alterations in actionable-druggable genes. We required that all amino acid sequence changes have either a corrected p-value < 0.05 and usage change of > 10% or a corrected p-value < 0.001 and usage change of > 5%. We further required that if the sequence alteration was dependent on a variant, the variant had to have a GENIE PolyPhen rating of probably_damaging to be reported. For gene fusions, we required that the fusion be present at at least 20% of the actionable druggable gene expression in the tumor and less than 10% of the gene expression in the normal sample.

### Generation of simulated data for testing variant calling

We first selected two annotated transcripts (GENCODE v38) each from 100 random genes on human chromosome 22, requiring that both transcripts contained at least one intron. We then randomly inserted two variants into each transcript, with 50% of transcripts having one SNV and one indel and the other 50% having two indels. The indels were constrained to be between 1 and 25bp long and the insertions contained random sequence. SNVs were constructed by randomly selecting one position, then changing the base at that position to another random distinct base. We generated a transcriptome fasta containing the variants, as well as a reference VCF file with variant positions. We then used Badread (v0.4.1)[70] to simulate 10X coverage of this transcriptome using the PacBio 2021 error model and otherwise default options.

### Evaluation of variant calling

For evaluating variant calling, we ran the following tools:

Longshot v1.0.0 with: longshot --force_overwrite --bam SAMPLE.bam --ref GENOME.fa --out SAMPLE.vcf --min_mapq 10 --strand_bias_pvalue_cutoff 0.000001

Clair3 v1.0.10 with: run_clair3.sh --bam_fn=SAMPLE.bam --ref_fn=GENOME.fa --threads=12 --platform=“hifi” --model_path=clair3/bin/models/hifi_sequel2 --output=SAMPLE

DeepVariant v1.8.0 through singularity with: run_deepvariant --model_type=PACBIO --ref=GENOME.fa --reads=SAMPLE.bam --output_vcf=SAMPLE.vcf.gz

--output_gvcf=SAMPLE.g.vcf.gz --num_shards=4

longcallR v1.12.0 with: longcallR --bam-path SAMPLE.bam --ref-path GENOME.fa --output SAMPLE.vcf --threads 8 --preset hifi-masseq

Clair3-RNA v0.2.2 with: /path/to/run_clari3_rna --bam_fn SAMPLE.bam --ref_fn GENOME.fa --threads 12 --platform hifi_sequel2_minimap2 –output_dir /path/to/output

To try to improve precision and recall for Longshot, we tested a CIGAR-splitting approach[71]:

1. gatk v4.6.2.0 with: gatk CreateSequenceDictionary -R GENOME.fa
2. gatk v4.6.2.0 with: gatk SplitNCigarReads -R GENOME.fa -I SAMPLE.bam -O SAMPLE.cigarSplit.bam
3. Longshot v1.0.0 with: longshot --bam SAMPLE.cigarSplit.bam --ref GENOME.fa --min_cov 3 --min_alt_count 3 -q 0 --max_cigar_indel 1000000 --out SAMPLE.vcf

We also obtained genomic variant calls (from Hi-C phased genotyping) for the same cell lines from ENCODE (MCF-10A: ENCFF354WRB, MCF-7: ENCFF216DQP, PC-9: ENCFF571MXE)[41,42].

We then filtered both the genomic and RNA variants to those that overlapped with expressed exons from running FLAIR collapse on the samples and had read coverage >20 reads. We then calculated recall of the RNA variants detected by each tool when compared to the genomic variants.

### Simulated fusion dataset

A fusion benchmarking dataset was originally designed by Haas et al, but we used the exact simulated reads of this dataset from the JAFFAL paper[24]. As in that paper, we combined the simulated fusion reads with NA12878 long-read ONT data from Workman et al[72] as background.

### Generating additional simulated fusion data for benchmarking

#### Fusions with non-genic regions

We first parsed GENCODE v38 annotation[63], keeping only those with at least one transcript with two or more exons. We then randomly selected 100 genes, randomly selecting breakpoints but ensuring that the breakpoints were after the first splice junction and before the last splice junction of at least one transcript. We then randomly selected one transcript and extracted the genomic sequence for the exons that were 5’ relative to the breakpoint. We next randomly selected a chromosome position, strand, and length, extracted the genomic sequence from hg38 (requiring that it contained no N bases). This formed the 3’ end of the fusion. We generated a FASTA and BED file representing the true fusion sequence model.

#### 3-gene fusions

We first parsed GENCODE v38 annotation[63], keeping only those with at least one transcript with three or more exons. For 100 iterations, we then randomly selected sets of three genes and randomly selected one isoform of each gene. For each isoform, we selected a random number of exons (X) up to the number of exons in the isoform (N) minus two. For the 5’ gene, we selected the first X exons, for the middle gene, we selected a random start position between exon two and exon N-X, and for the 3’ gene, we selected the last X exons. We then extracted the genomic sequence of those exons and generated a FASTA and BED file representing the true fusion sequence model.

#### Fusion isoforms

We first parsed GENCODE v38 annotation[63], keeping only those with at least two transcripts with two or more exons. For 100 iterations, we then randomly selected sets of two genes and randomly selected a breakpoint position for each gene, ensuring that the breakpoints were after the first splice junction and before the last splice junction of at least two transcripts. We then randomly selected two isoforms for each gene. We randomly paired the isoforms and assigned one gene to be 5’ and the other 3’, then extracted the genomic sequence of the exons and generated a FASTA and BED file representing the true fusion sequence models.

#### Fusion simulation

All simulated datasets in this section were simulated with Badread (v0.4.1)[70] with the PacBio preset function, which is:

badread simulate --reference ref.fasta --quantity Nx --error_model pacbio2021 --qscore_model pacbio2021 --identity 30,3 > reads.fastq.gz

We simulated all datasets at the coverage levels (N) 5, 10, 50, and 100x.

### Comparison to other gene fusion calling tools

CTAT-LR-fusion v0.13.0[25] was run using docker and using the GRCh38_gencode_v37_CTAT_lib_Mar012021.plug-n-play reference. We then converted gene names to be compatible with GENCODE v38 based on genomic coordinates.

JAFFAL v2.3[24] was run with <path to JAFFA>/tools/bin/bpipe run <path to JAFFA>/JAFFAL.groovy fastq.gz. This uses their default genome and annotation which are hg38 and GENCODE v22. We also converted gene names from the JAFFAL output to GENCODE v38 compatible names.

LongGF[43] was run with reads aligned to the hg38 genome and the GENCODE v38 reference gtf and default settings: LongGF sorted.bam gencode.v37.annotation.gtf 100 50 200.

The output of all tools were parsed and fusions extracted by gene/locus names. Fusions were considered matching if both gene names matched.

### KEAP1 protein structure comparison

We used AlphaFold[73] to generate a predicted protein folding structure for the KEAP1 sequence with the 10AA deletion. We then compared that to the normal predicted KEAP1 structure from AlphaFold[74] using the Protein Data Bank Pairwise Structure Alignment tool TM-align method[75].

### Comparison with short-read RNA-seq of osteosarcoma PDX-derived cell lines

Short-reads for these same cell lines were obtained from Schott et al[45]. These were aligned with STAR v2.7.11b[76] and gene counts were identified using featureCounts[77].

### WGS structural variant and SNV detection

These data were obtained as described in Schott et al[45].

## Declarations

### Ethics approval and consent to participate

Osteosarcoma samples are previously published in Sayles et al[44] and were consented as part of that study.

For studies related to samples FH-1427 and FH-1633, patients were consented on tissue collection, tumor sequencing, and sharing of deidentified data in restricted-access databases, with approval by the University of Washington Institutional Review Board (Protocol Number RG1000720).

### Consent for publication

Not applicable

### Availability of data and materials

The datasets generated and/or analysed during the current study will be available in dbGaP. All code for doing novel fusion and fusion isoform simulations is available at https://github.com/cafelton/testfusionisoforms. FLAIR3 code is available at https://github.com/BrooksLabUCSC/flair

## Authors’ contributions

**Colette Felton** conceptualization, methodology, software, validation, formal analysis, investigation, data curation, writing, visualization

Andrea Galvez investigation

Tanvi Damle validation, formal analysis, visualization

Kevin M. Levine resources

Mark Diekhans software

Eunice Lopez Fuentes validation

Taylor S. Won validation

Christopher Vollmers methodology, investigation

Alejandro Sweet-Cordero resources, supervision

Alice H. Berger resources, supervision, funding acquisition

**Angela N Brooks** conceptualization, methodology, writing - review, supervision, project administration, funding acquisition

## Acknowledgements

PacBio sequencing data were collected at the University of Washington Long Reads Sequencing Center. We thank Pamela Yang at the Fred Hutch Cancer Center Clinical Research Specimen Processing Shared Resource for assistance with RNA extraction. We also thank Leanne Sayles, Marcus Breese, Andrew Clugston, and Alex Lee at UCSF for their assistance with the osteosarcoma data and samples.

## Supplementary Legends

**Supplementary File 1: FLAIR3 isoform calls for osteosarcoma samples.** This is a bed format file with CDS information included. For the name field, the format is: [Splice junction chain ID]-[unique ends ID]_[transcript name]_[productivity]_[gene].

**Supplementary File 2: FLAIR3 isoform counts for osteosarcoma samples.** Column 1: Isoform IDs in format [Splice junction chain ID]-[unique ends ID]_[transcript name]_[gene]. Columns 2-13: read counts for each replicate of each cell line.

**Supplementary File 3: FLAIR3 isoform calls for lung samples.** This is a bed format file with CDS information included. For the name field, the format is: [Splice junction chain ID]-[unique ends ID]_[transcript name]_[productivity]_[gene].

**Supplementary File 4: FLAIR3 isoform counts for lung samples.** Column 1: Isoform IDs in format [Splice junction chain ID]-[unique ends ID]_[transcript name]_[gene]. Columns 2-5: read counts for each sample.

## References

1. Gibbs SN, Peneva D, Cuyun Carter G, Palomares MR, Thakkar S, Hall DW, et al. Comprehensive review on the clinical impact of next-generation sequencing tests for the management of advanced cancer. JCO Precis Oncol. American Society of Clinical Oncology; 2023;7:e2200715.

2. Colomer R, Miranda J, Romero-Laorden N, Hornedo J, González-Cortijo L, Mouron S, et al. Usefulness and real-world outcomes of next generation sequencing testing in patients with cancer: an observational study on the impact of selection based on clinical judgement. EClinicalMedicine. Elsevier BV; 2023;60:102029.

3. Menzel M, Ossowski S, Kral S, Metzger P, Horak P, Marienfeld R, et al. Multicentric pilot study to standardize clinical whole exome sequencing (WES) for cancer patients. NPJ Precis Oncol. Springer Science and Business Media LLC; 2023;7:106.

4. Xiao W, Ren L, Chen Z, Fang LT, Zhao Y, Lack J, et al. Toward best practice in cancer mutation detection with whole-genome and whole-exome sequencing. Nat Biotechnol. Springer Science and Business Media LLC; 2021;39:1141–50.

5. Hong M, Tao S, Zhang L, Diao L-T, Huang X, Huang S, et al. RNA sequencing: new technologies and applications in cancer research. J Hematol Oncol. Springer Science and Business Media LLC; 2020;13:166.

6. Pardo-Palacios FJ, Wang D, Reese F, Diekhans M, Carbonell-Sala S, Williams B, et al. Systematic assessment of long-read RNA-seq methods for transcript identification and quantification [Internet]. bioRxiv. 2023 [cited 2023 July 30]. p. 2023.07.25.550582. 10.1101/2023.07.25.550582

7. Soulette CM, Hrabeta-Robinson E, Arevalo C, Felton C, Tang AD, Marin MG, et al. Full-length transcript alterations in human bronchial epithelial cells with U2AF1 S34F mutations. Life Sci Alliance [Internet]. 2023;6. 10.26508/lsa.202000641

8. Tang AD, Soulette CM, van Baren MJ, Hart K, Hrabeta-Robinson E, Wu CJ, et al. Full-length transcript characterization of SF3B1 mutation in chronic lymphocytic leukemia reveals downregulation of retained introns. Nat Commun. England; 2020;11:1438.

9. Edge P, Bansal V. Longshot enables accurate variant calling in diploid genomes from single-molecule long read sequencing. Nat Commun. 2019;10:4660.

10. Volden R, Palmer T, Byrne A, Cole C, Schmitz RJ, Green RE, et al. Improving nanopore read accuracy with the R2C2 method enables the sequencing of highly multiplexed full-length single-cell cDNA. Proc Natl Acad Sci U S A. 2018;115:9726–31.

11. Adams M, Vollmers C. Generation and analysis of a mouse multi-tissue genome annotation atlas. bioRxivorg. 2024;2024.01.31.578267.

12. Ni Y, Liu X, Simeneh ZM, Yang M, Li R. Benchmarking of Nanopore R10.4 and R9.4.1 flow cells in single-cell whole-genome amplification and whole-genome shotgun sequencing. Comput Struct Biotechnol J. Elsevier BV; 2023;21:2352–64.

13. Shi Z-X, Chen Z-C, Zhong J-Y, Hu K-H, Zheng Y-F, Chen Y, et al. High-throughput and high-accuracy single-cell RNA isoform analysis using PacBio circular consensus sequencing. Nat Commun. Springer Science and Business Media LLC; 2023;14:2631.

14. Shafin K, Pesout T, Chang P-C, Nattestad M, Kolesnikov A, Goel S, et al. Haplotype-aware variant calling with PEPPER-Margin-DeepVariant enables high accuracy in nanopore long-reads. Nat Methods. Springer Science and Business Media LLC; 2021;18:1322–32.

15. Ahsan MU, Liu Q, Fang L, Wang K. NanoCaller for accurate detection of SNPs and indels in difficult-to-map regions from long-read sequencing by haplotype-aware deep neural networks. Genome Biol. Springer Science and Business Media LLC; 2021;22:261.

16. Mullaney JM, Mills RE, Pittard WS, Devine SE. Small insertions and deletions (INDELs) in human genomes. Hum Mol Genet. Oxford University Press (OUP); 2010;19:R131–6.

17. Haas BJ, Dobin A, Li B, Stransky N, Pochet N, Regev A. Accuracy assessment of fusion transcript detection via read-mapping and de novo fusion transcript assembly-based methods. Genome Biol. 2019;20:213.

18. Lin J, Jia P, Wang S, Kosters W, Ye K. Comparison and benchmark of structural variants detected from long read and long-read assembly. Brief Bioinform. Oxford University Press (OUP); 2023;24:bbad188.

19. Mahmoud M, Gobet N, Cruz-Dávalos DI, Mounier N, Dessimoz C, Sedlazeck FJ. Structural variant calling: the long and the short of it. Genome Biol. Springer Science and Business Media LLC; 2019;20:246.

20. Felton C, Tang AD, Knisbacher B, Wu CJ, Brooks AN. Detection of alternative isoforms of gene fusions from long-read RNA-seq with FLAIR-fusion [Internet]. bioRxiv. 2022 [cited 2022 Dec 8]. p. 2022.08.01.502364. 10.1101/2022.08.01.502364

21. Mohammad T, Zolotovskaia MA, Suntsova MV, Buzdin AA. Cancer fusion transcripts with human non-coding RNAs. Front Oncol. Frontiers Media SA; 2024;14:1415801.

22. Zhong X, Luan J, Yu A, Lee-Hassett A, Miao Y, Yang L. SFyNCS detects oncogenic fusions involving non-coding sequences in cancer. Nucleic Acids Res. Oxford University Press (OUP); 2023;51:e96.

23. PCAWG Transcriptome Core Group, Calabrese C, Davidson NR, Demircioğlu D, Fonseca NA, He Y, et al. Genomic basis for RNA alterations in cancer. Nature. 2020;578:129–36.

24. Davidson NM, Chen Y, Sadras T, Ryland GL, Blombery P, Ekert PG, et al. JAFFAL: detecting fusion genes with long-read transcriptome sequencing. Genome Biol. Springer Science and Business Media LLC; 2022;23:10.

25. Qin Q, Popic V, Yu H, White E, Khorgade A, Shin A, et al. CTAT-LR-fusion: accurate fusion transcript identification from long and short read isoform sequencing at bulk or single cell resolution. bioRxivorg. 2024;2024.02.24.581862.

26. Robles-Espinoza CD, Mohammadi P, Bonilla X, Gutierrez-Arcelus M. Allele-specific expression: applications in cancer and technical considerations. Curr Opin Genet Dev. 2021;66:10–9.

27. Durham M, Vadde S, Brooks AN. MET exon 14 skipping is overexpressed in an allele-specific manner in lung adenocarcinoma primary samples. MicroPubl Biol [Internet]. California Institute of Technology; 2023 [cited 2024 Aug 16];2023. 10.17912/micropub.biology.000957

28. Esfahani MS, Lee LJ, Jeon Y-J, Flynn RA, Stehr H, Hui AB, et al. Functional significance of U2AF1 S34F mutations in lung adenocarcinomas. Nat Commun. 2019;10:5712.

29. Tang AD, Hrabeta-Robinson E, Volden R, Vollmers C, Brooks AN. Detecting haplotype-specific transcript variation in long reads with FLAIR2. bioRxiv [Internet]. 2023; 10.1101/2023.06.09.544396

30. Glinos DA, Garborcauskas G, Hoffman P, Ehsan N, Jiang L, Gokden A, et al. Transcriptome variation in human tissues revealed by long-read sequencing. Nature. Springer Science and Business Media LLC; 2022;608:353–9.

31. Quinones-Valdez G, Amoah K, Xiao X. Long-read RNA-seq demarcates cis- and trans-directed alternative RNA splicing. Nat Commun. Springer Science and Business Media LLC; 2025;16:9603.

32. Byrne A, Le D, Sereti K, Menon H, Vaidya S, Patel N, et al. Single-cell long-read targeted sequencing reveals transcriptional variation in ovarian cancer. Nat Commun. Springer Science and Business Media LLC; 2024;15:6916.

33. Miller RM, Jordan BT, Mehlferber MM, Jeffery ED, Chatzipantsiou C, Kaur S, et al. Enhanced protein isoform characterization through long-read proteogenomics. Genome Biol. Springer Science and Business Media LLC; 2022;23:69.

34. Veiga DFT, Nesta A, Zhao Y, Deslattes Mays A, Huynh R, Rossi R, et al. A comprehensive long-read isoform analysis platform and sequencing resource for breast cancer. Sci Adv. 2022;8:eabg6711.

35. Ishino T, Watanabe T, Tokita S, Ueda Y, Kawase K, Takano Y, et al. Immunopeptidomics combined with full-length transcriptomics uncovers diverse neoantigens. Cell Rep. Elsevier BV; 2026;45:116781.

36. Jetzinger F, Paniagua A, Cormack S, Mestre-Tomás J, Morante-Redolat JM, Fariñas I, et al. Handling biological replicates in long-read RNA sequencing data by joining or not joining. Nat Commun [Internet]. Springer Science and Business Media LLC; 2026;17. 10.1038/s41467-026-77367-6

37. Tang AD, Felton C, Hrabeta-Robinson E, Volden R, Vollmers C, Brooks AN. Detecting haplotype-specific transcript variation in long reads with FLAIR2. Genome Biol. Springer Science and Business Media LLC; 2024;25:173.

38. Zheng Z, Li S, Su J, Leung AW-S, Lam T-W, Luo R. Symphonizing pileup and full-alignment for deep learning-based long-read variant calling. Nat Comput Sci. Springer Science and Business Media LLC; 2022;2:797–803.

39. Zheng Z, Yu X, Chen L, Lee Y-L, Xin C, Wong AOK, et al. Clair3-RNA: a deep learning-based small variant caller for long-read RNA sequencing data. Nat Commun. Springer Science and Business Media LLC; 2025;16:11553.

40. Huang N, Li H, Human Pangenome Reference Consortium. SNP calling, haplotype phasing and allele-specific analysis with long RNA-seq reads. Nat Methods. Springer Science and Business Media LLC; 2026;23:1962–7.

41. Luo Y, Hitz BC, Gabdank I, Hilton JA, Kagda MS, Lam B, et al. New developments on the Encyclopedia of DNA Elements (ENCODE) data portal. Nucleic Acids Res. Oxford University Press (OUP); 2020;48:D882–9.

42. ENCODE Project Consortium. An integrated encyclopedia of DNA elements in the human genome. Nature. 2012;489:57–74.

43. Liu Q, Hu Y, Stucky A, Fang L, Zhong JF, Wang K. LongGF: computational algorithm and software tool for fast and accurate detection of gene fusions by long-read transcriptome sequencing. BMC Genomics. 2020;21:793.

44. Sayles LC, Breese MR, Koehne AL, Leung SG, Lee AG, Liu H-Y, et al. Genome-Informed Targeted Therapy for Osteosarcoma. Cancer Discov. 2019;9:46–63.

45. Schott CR, Koehne AL, Sayles LC, Young EP, Luck C, Yu K, et al. Osteosarcoma PDX-derived cell line models for preclinical drug evaluation demonstrate metastasis inhibition by dinaciclib through a genome-targeted approach. Clin Cancer Res. American Association for Cancer Research (AACR); 2024;30:849–64.

46. Chen X, Bahrami A, Pappo A, Easton J, Dalton J, Hedlund E, et al. Recurrent somatic structural variations contribute to tumorigenesis in pediatric osteosarcoma. Cell Rep. Elsevier BV; 2014;7:104–12.

47. Zhou B, Lin W, Long Y, Yang Y, Zhang H, Wu K, et al. Notch signaling pathway: architecture, disease, and therapeutics. Signal Transduct Target Ther. Springer Science and Business Media LLC; 2022;7:95.

48. Zavitsanou A-M, Pillai R, Hao Y, Wu WL, Bartnicki E, Karakousi T, et al. KEAP1 mutation in lung adenocarcinoma promotes immune evasion and immunotherapy resistance. Cell Rep. Elsevier BV; 2023;42:113295.

49. Morgenstern C, Lastres-Becker I, Demirdöğen BC, Costa VM, Daiber A, Foresti R, et al. Biomarkers of NRF2 signalling: Current status and future challenges. Redox Biol. Elsevier BV; 2024;72:103134.

50. Chalmers ZR, Connelly CF, Fabrizio D, Gay L, Ali SM, Ennis R, et al. Analysis of 100,000 human cancer genomes reveals the landscape of tumor mutational burden. Genome Med. BioMed Central; 2017;9:34.

51. Li H, Pan Y, Li Y, Li C, Wang R, Hu H, et al. Frequency of well-identified oncogenic driver mutations in lung adenocarcinoma of smokers varies with histological subtypes and graduated smoking dose. Lung Cancer. Elsevier BV; 2013;79:8–13.

52. Cancer Genome Atlas Research Network. Comprehensive molecular profiling of lung adenocarcinoma. Nature. 2014;511:543–50.

53. Hagen RM, Adamo P, Karamat S, Oxley J, Aning JJ, Gillatt D, et al. Quantitative analysis of ERG expression and its splice isoforms in formalin-fixed, paraffin-embedded prostate cancer samples: association with seminal vesicle invasion and biochemical recurrence. Am J Clin Pathol. Oxford University Press (OUP); 2014;142:533–40.

54. East P, Kelly GP, Biswas D, Marani M, Hancock DC, Creasy T, et al. RAS oncogenic activity predicts response to chemotherapy and outcome in lung adenocarcinoma. Nat Commun. Springer Science and Business Media LLC; 2022;13:5632.

55. AACR Project GENIE Consortium. AACR Project GENIE: Powering precision medicine through an international consortium. Cancer Discov. 2017;7:818–31.

56. Cerami E, Gao J, Dogrusoz U, Gross BE, Sumer SO, Aksoy BA, et al. The cBio cancer genomics portal: an open platform for exploring multidimensional cancer genomics data. Cancer Discov. 2012;2:401–4.

57. Joruiz SM, Beck JA, Horikawa I, Harris CC. The Δ133p53 isoforms, tuners of the p53 pathway. Cancers (Basel). MDPI AG; 2020;12:3422.

58. Yang R, Serrano M, Slater J, Leung E, Koeffler HP. Analysis of p16INK4a and its interaction with CDK4. Biochem Biophys Res Commun. Elsevier BV; 1996;218:254–9.

59. Stott FJ, Bates S, James MC, McConnell BB, Starborg M, Brookes S, et al. The alternative product from the human CDKN2A locus, p14(ARF), participates in a regulatory feedback loop with p53 and MDM2. EMBO J. Wiley; 1998;17:5001–14.

60. Podda MS, Tatoni D, Mattei G, Magi A, D’Aurizio R, Poliseno L. Landscape of BRAF transcript variants in human cancer. Mol Oncol. Mol Oncol; 2025;19:2700–14.

61. Klonowski J, Liang Q, Coban-Akdemir Z, Lo C, Kostka D. Aenmd: Annotating escape from nonsense-mediated decay for transcripts with protein-truncating variants. Bioinformatics. Oxford University Press (OUP); 2023;39:btad556.

62. Danecek P, Bonfield JK, Liddle J, Marshall J, Ohan V, Pollard MO, et al. Twelve years of SAMtools and BCFtools. Gigascience [Internet]. 2021;10. 10.1093/gigascience/giab008

63. Frankish A, Diekhans M, Jungreis I, Lagarde J, Loveland JE, Mudge JM, et al. GENCODE 2021. Nucleic Acids Res. 2021;49:D916–23.

64. Love MI, Huber W, Anders S. Moderated estimation of fold change and dispersion for RNA-seq data with DESeq2. Genome Biol. 2014;15:550.

65. Li H. Minimap2: pairwise alignment for nucleotide sequences. Bioinformatics. 2018;34:3094–100.

66. Adzhubei I, Jordan DM, Sunyaev SR. Predicting functional effect of human missense mutations using PolyPhen-2. Curr Protoc Hum Genet. Wiley; 2013;Chapter 7:Unit7.20.

67. Nagy E, Maquat LE. A rule for termination-codon position within intron-containing genes: when nonsense affects RNA abundance. Trends Biochem Sci. Elsevier BV; 1998;23:198–9.

68. UniProt Consortium. UniProt: The universal protein knowledgebase in 2025. Nucleic Acids Res. Oxford University Press (OUP); 2025;53:D609–17.

69. Nowicka M, Robinson MD. DRIMSeq: a Dirichlet-multinomial framework for multivariate count outcomes in genomics. F1000Res. 2016;5:1356.

70. Wick R. Badread: simulation of error-prone long reads. J Open Source Softw. The Open Journal; 2019;4:1316.

71. de Souza VBC, Jordan BT, Tseng E, Nelson EA, Hirschi KK, Sheynkman G, et al. Transformation of alignment files improves performance of variant callers for long-read RNA sequencing data. Genome Biol. 2023;24:91.

72. Workman RE, Tang AD, Tang PS, Jain M, Tyson JR, Razaghi R, et al. Nanopore native RNA sequencing of a human poly(A) transcriptome. Nat Methods. United States; 2019;16:1297–305.

73. Jumper J, Evans R, Pritzel A, Green T, Figurnov M, Ronneberger O, et al. Highly accurate protein structure prediction with AlphaFold. Nature. Springer Science and Business Media LLC; 2021;596:583–9.

74. Varadi M, Bertoni D, Magana P, Paramval U, Pidruchna I, Radhakrishnan M, et al. AlphaFold Protein Structure Database in 2024: providing structure coverage for over 214 million protein sequences. Nucleic Acids Res. Oxford University Press (OUP); 2024;52:D368–75.

75. Bittrich S, Segura J, Duarte JM, Burley SK, Rose Y. RCSB protein Data Bank: exploring protein 3D similarities via comprehensive structural alignments. Bioinformatics. Oxford University Press (OUP); 2024;40:btae370.

76. Dobin A, Davis CA, Schlesinger F, Drenkow J, Zaleski C, Jha S, et al. STAR: ultrafast universal RNA-seq aligner. Bioinformatics. Oxford University Press (OUP); 2013;29:15–21.

77. Liao Y, Smyth GK, Shi W. featureCounts: an efficient general purpose program for assigning sequence reads to genomic features. Bioinformatics. Oxford University Press (OUP); 2014;30:923–30.

