## Supplemental Figures for "Fully integrated SNV, fusion, and isoform analysis of cancer drivers with FLAIR3 for improved functional interpretation"

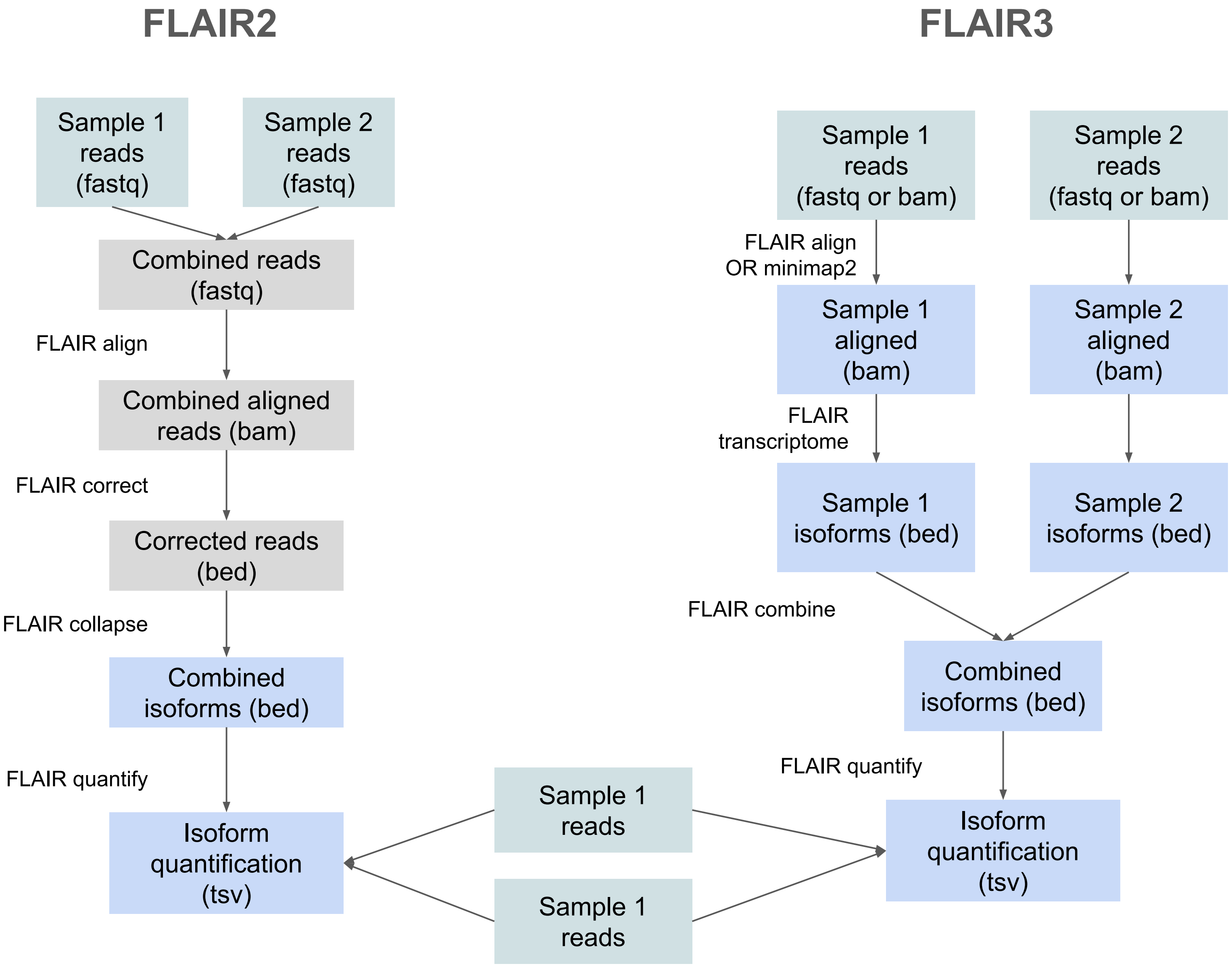


**Supplementary Figure 1: FLAIR analysis pipeline comparison between FLAIR2 and FLAIR3.** FLAIR3 introduces a method to run samples individually, then combine the transcriptomes at the end into a non-redundant set.

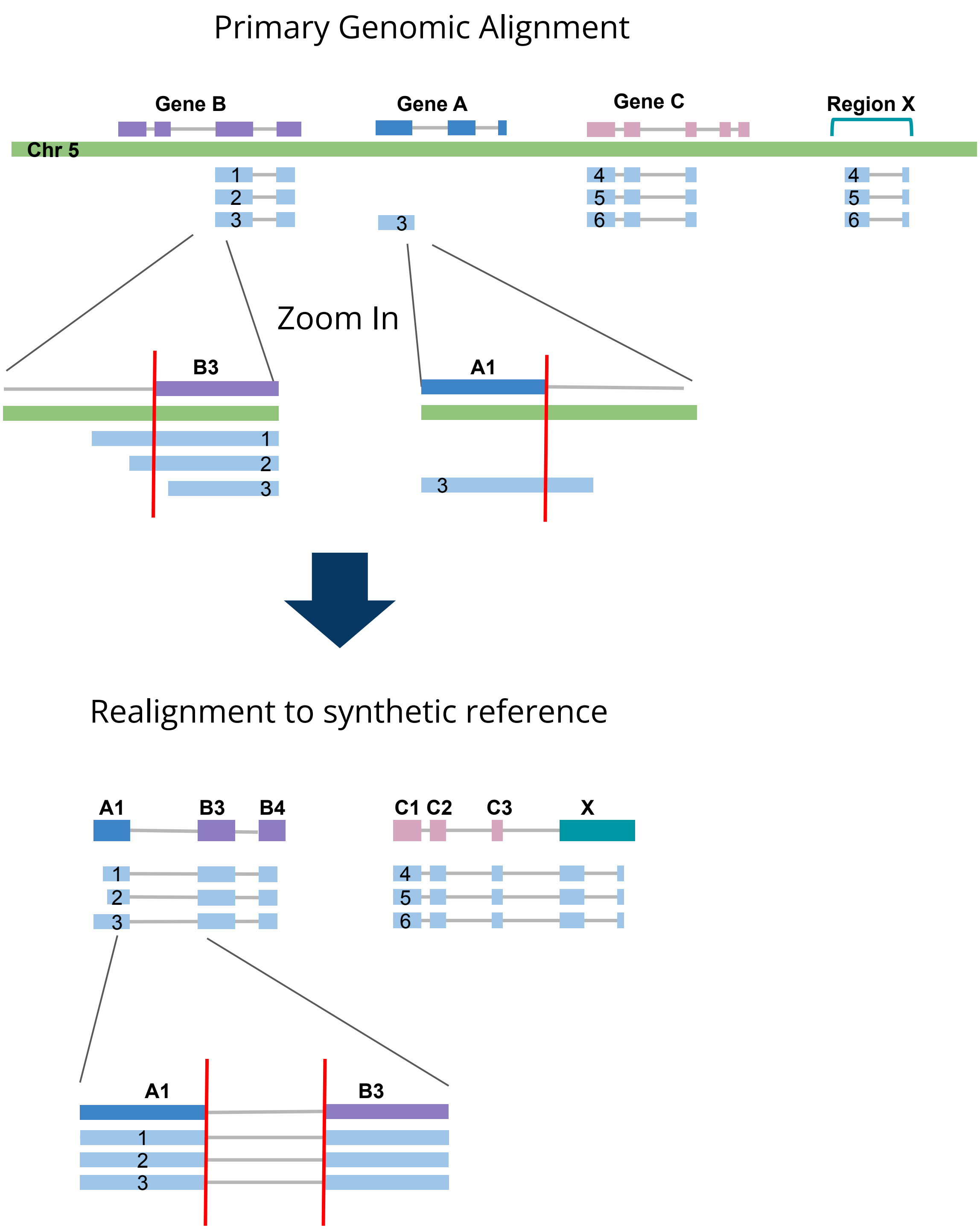


**Supplementary Figure 2: Schematic representation of how FLAIR fusion synthetic realignment improves breakpoint and quantification accuracy.** After genomic alignment, chimeric reads are identified through supplementary alignments. However, breakpoints may be messy due to double alignment of read sequence to both loci within the fusion. A synthetic fusion chromosome is constructed using sequence around the chimeric alignments, then reads are realigned to the model. The final fusion breakpoint is derived from realignment to the model.

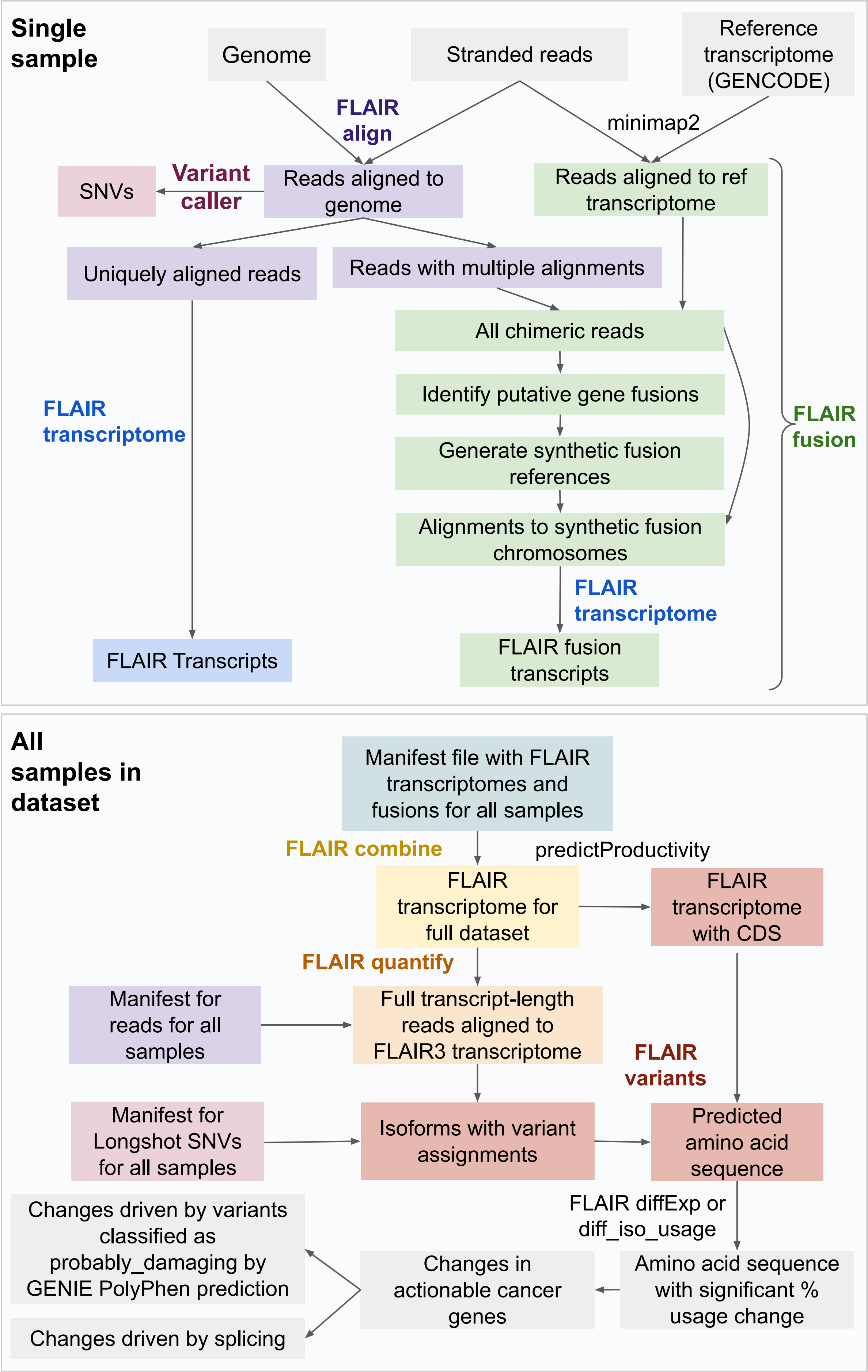


**Supplementary Figure 3: Expanded FLAIR3 pipeline including postprocessing.** This expanded pipeline shows how data flows within and between FLAIR3 steps (especially FLAIR fusion) and includes details on postprocessing results with GENIE annotation (see Methods).

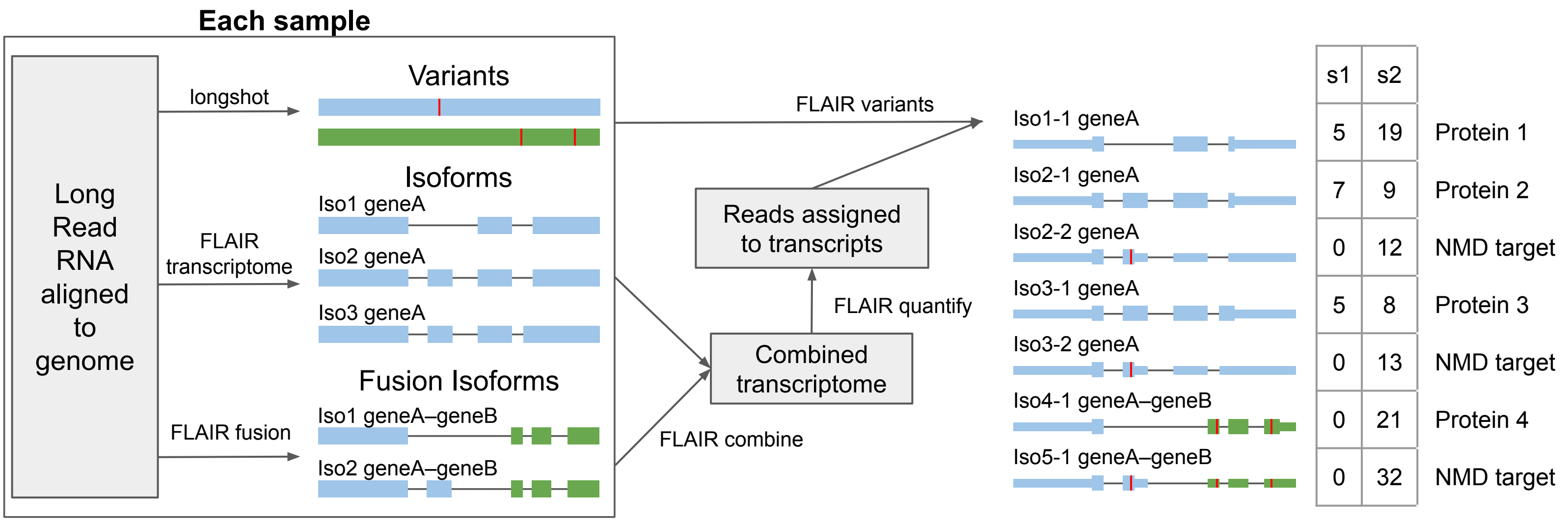


**Supplementary Figure 4: Integration of variants with transcripts and fusions.**

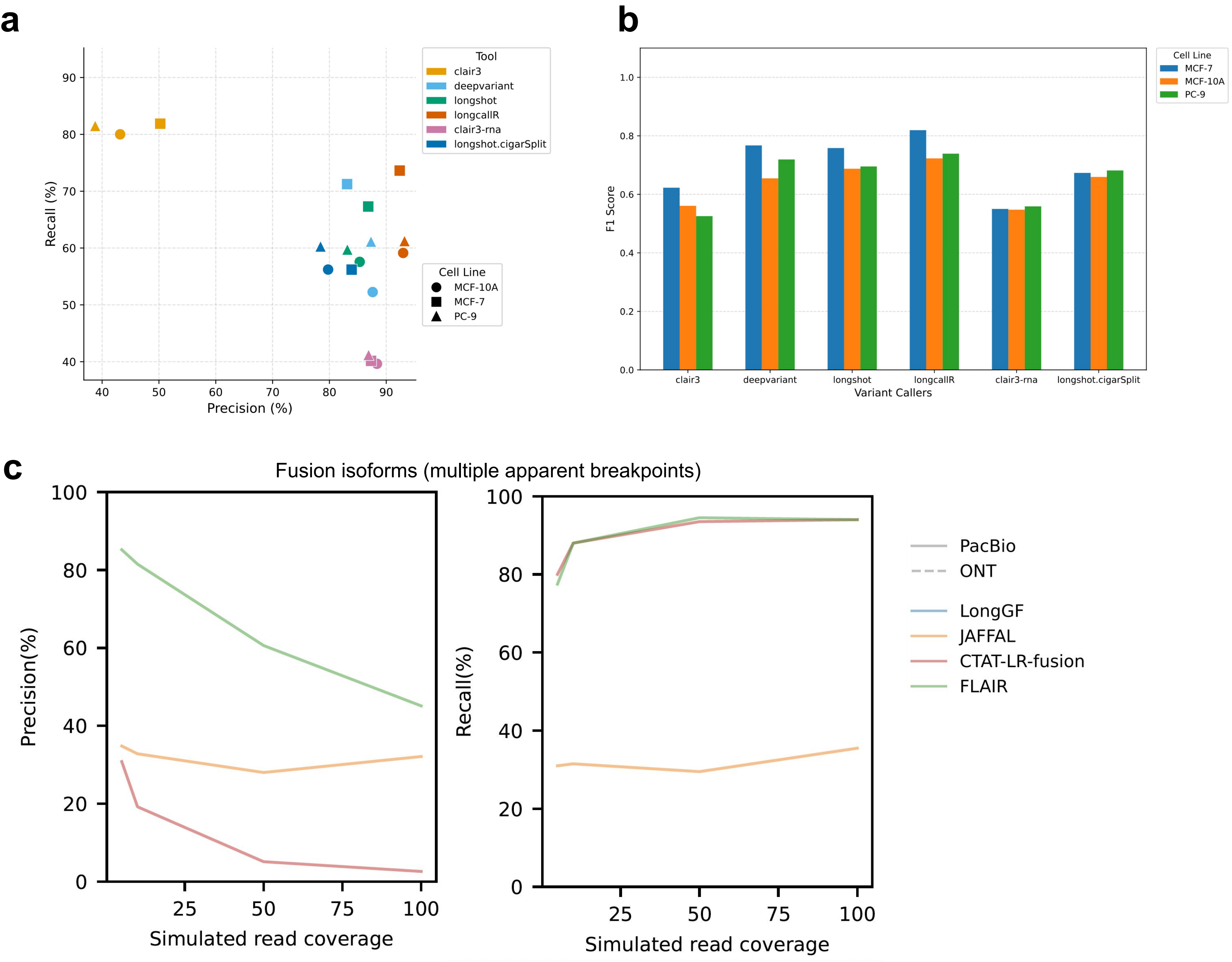


**Supplementary Figure 5: Performance of FLAIR3 fusion isoform detection and Longshot variant detection. a,** Precision and recall of Clair3, DeepVariant, Longshot, longcallR, Clari3-RNA, and Longshot + cigarSplit variant callers on SNVs in ENCODE PacBio sequencing of 3 cell lines. Calculated compared to matched DNA-sequencing of these cell lines. Variants called from RNA were only considered if they overlapped annotated exons and if positions had coverage of >10 reads. **c,** Bar plot of F1 scores for variant calling in 3 cell lines as in b. **c,** Recall (right) and precision (left) of fusion breakpoint identification (accuracy of within 1bp) on simulated fusions with multiple isoforms that create two apparent breakpoints.

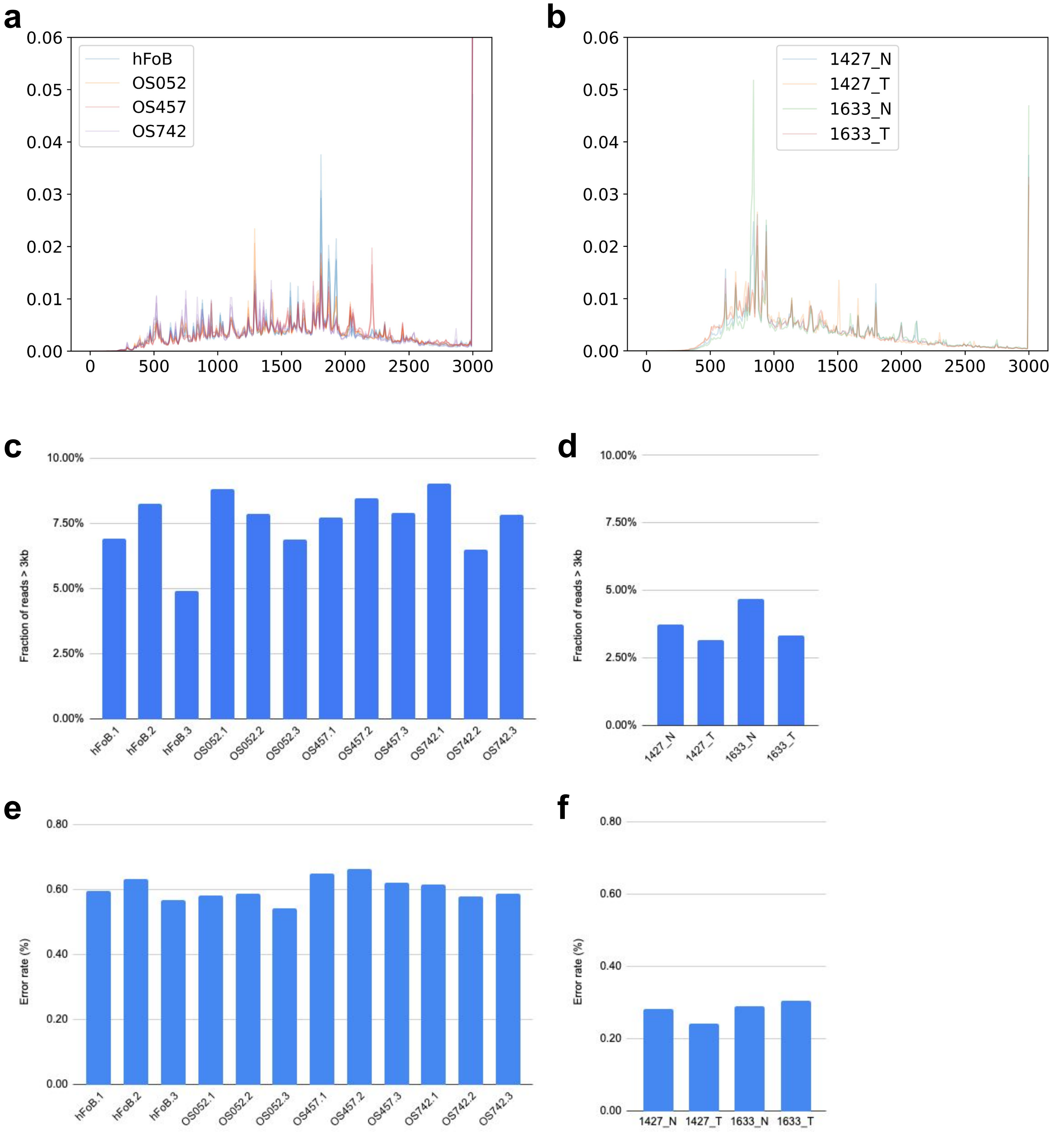


**Supplementary Figure 6: Length distribution and alignment accuracy of long-read RNA-seq data. a,** Length distribution of reads from osteosarcoma samples, generated with R2C2 + ONT. **b,** Length distribution of reads from lung samples, generated with PacBio. **c,** Fraction of reads >3kb from osteosarcoma samples. **d,** Fraction of reads >3kb from lung samples. **e,** Error rate in osteosarcoma samples. **f,** Error rate in lung samples.

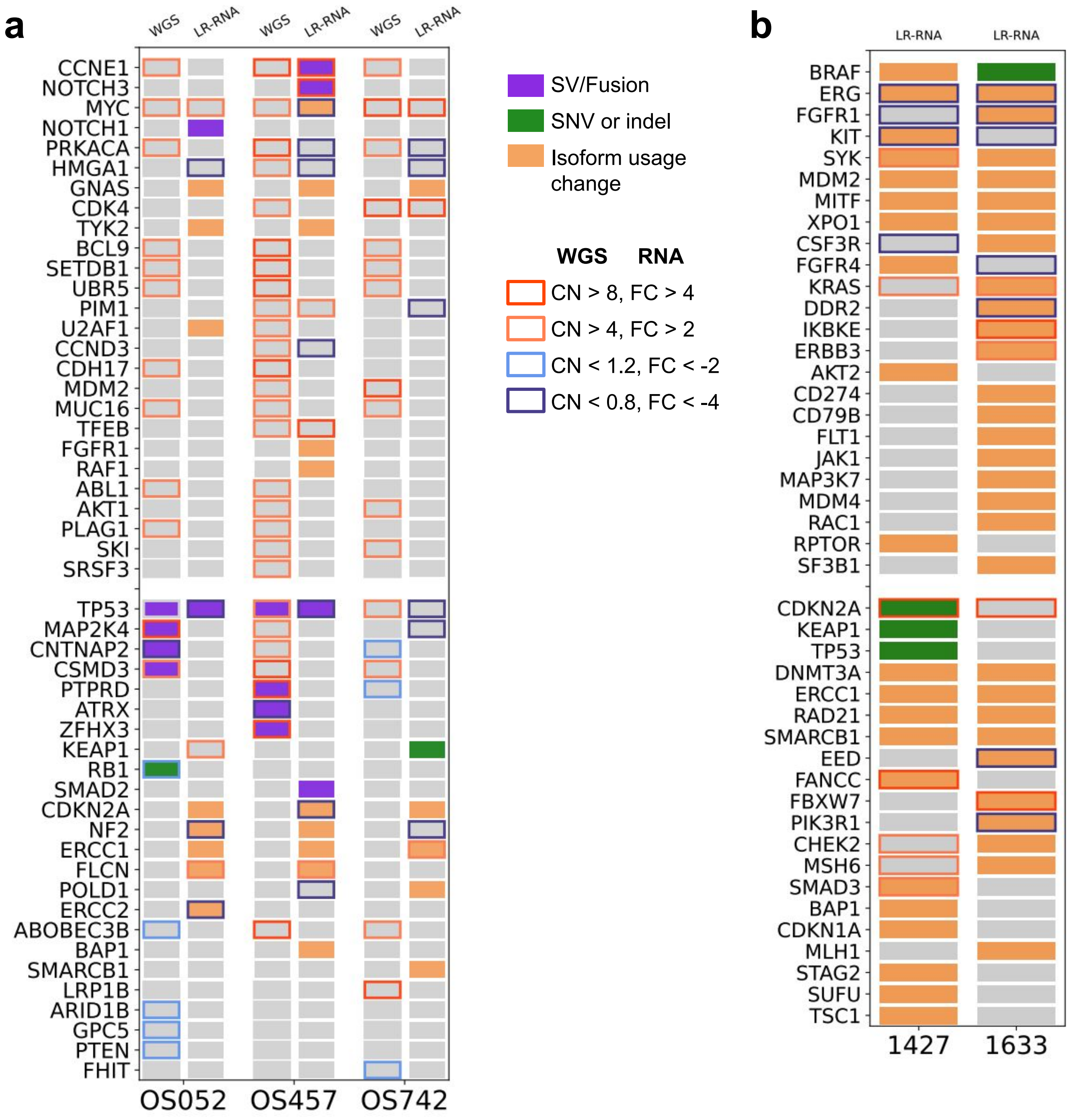


**Supplementary Figure 7: FLAIR3 identifies known and novel cancer drivers in multiple tumor types. a,** OncoPrint of cancer-relevant alterations in each osteosarcoma cell line. All statistics were done in comparison to the hFoB samples. Bottom x-axis shows the ID of each cell line, top x-axis shows the sequencing method, y-axis is genes (only actionable-druggable cancer genes included). Purple box = gene fusion, green box = SNV or indel, pale orange box = splicing change that causes change to the amino acid sequence produced by the gene (DRIMSeq p-val 0.05, delta usage > 10%). Borders: copy number (WGS, CN) or gene expression (lrRNA-seq) change (DESeq2 pval < 0.05). Dark blue: major loss or decrease , light blue: minor loss or decrease, orange: minor gain or increase, bright red: major gain or increase. **b,** OncoPrint of alterations in lung samples. Key is as described in b, but alterations identified by comparing the tumor and matched normal. Due to the lack of replicates, the gene expression changes were evaluated purely on fold change, while the splicing changes were calculated using a Fisher's exact test.

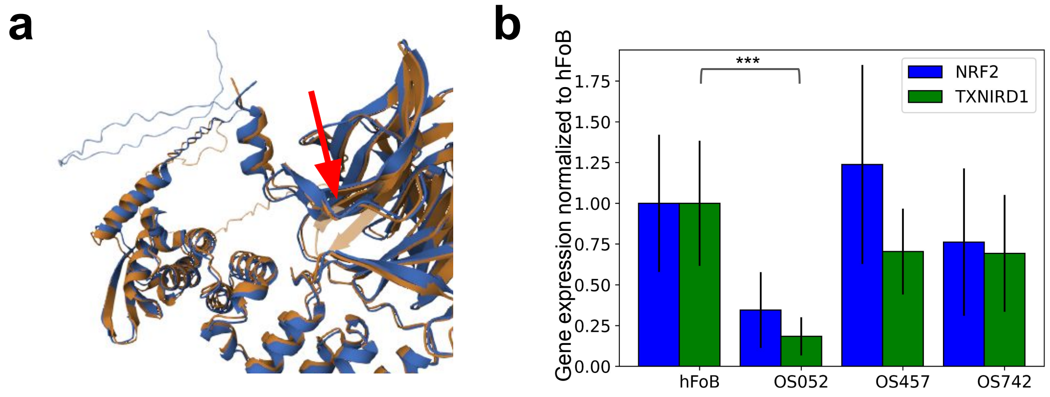


**Supplementary Figure 8: Structural and pathway analysis of 8AA deletion in *KEAP1* in osteosarcoma. a,** AlphaFold models of normal KEAP1 sequence (orange) and KEAP1 protein with deletion (blue). Altered region highlighted with red arrow. **b,** Bar plots of transcript-level expression of *NRF2*, which KEAP1 represses, and *TXNIRD1*, which NRF2 regulates.

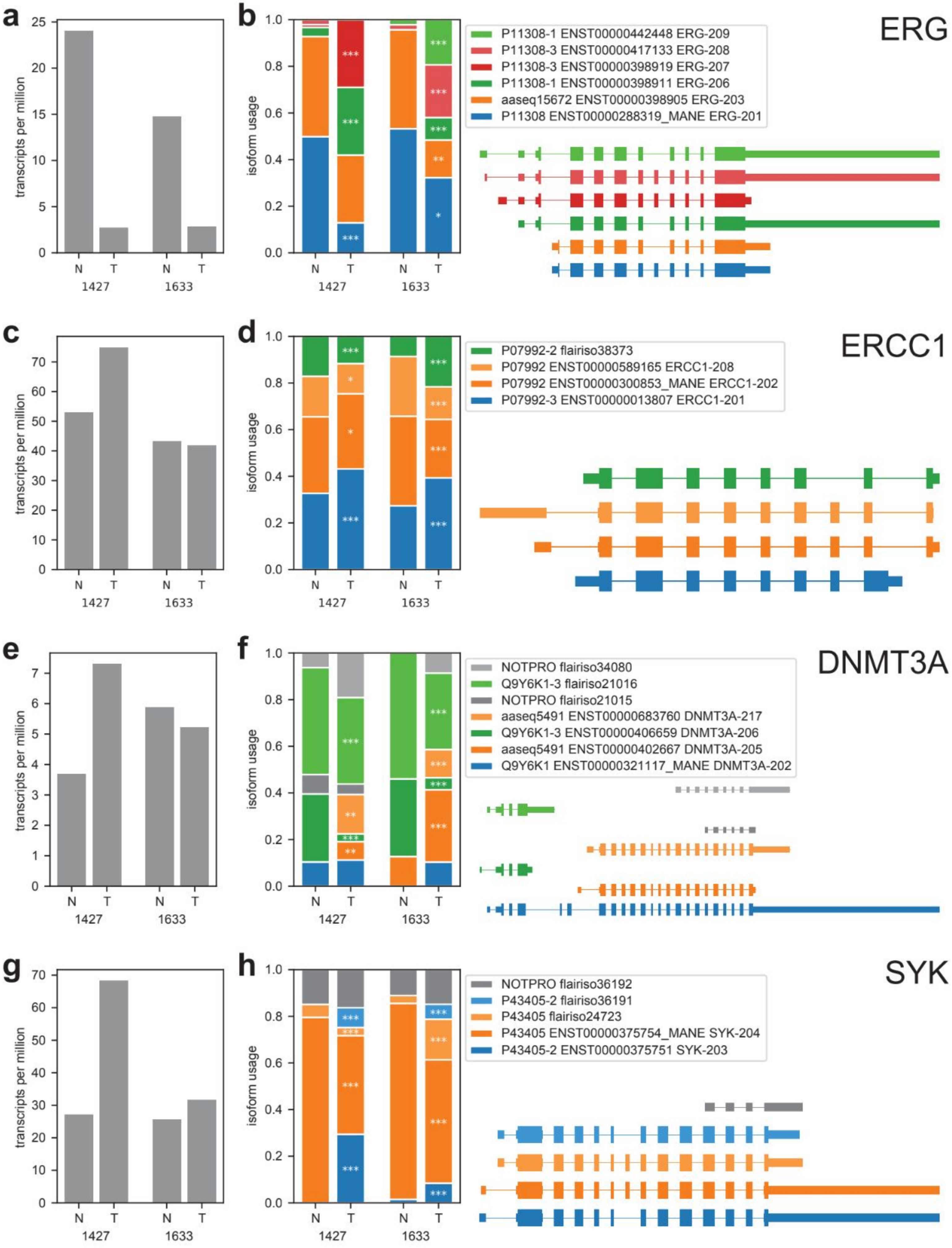
a
b
ERG
transcripts per million
isoform usage
P11308-1 ENST00000442448 ERG-209
P11308-3 ENST00000417133 ERG-208
P11308-3 ENST00000398919 ERG-207
P11308-1 ENST00000398911 ERG-206
aaseq15672 ENST00000398905 ERG-203
P11308 ENST00000288319_MANE ERG-201
N T N T
1427 1633
c
d
ERCC1
P07992-2 flairiso38373
P07992 ENST00000589165 ERCC1-208
P07992 ENST00000300853_MANE ERCC1-202
P07992-3 ENST00000013807 ERCC1-201
e
f
DNMT3A
NOTPRO flairiso34080
Q9Y6K1-3 flairiso21016
NOTPRO flairiso21015
aaseq5491 ENST00000683760 DNMT3A-217
Q9Y6K1-3 ENST00000406659 DNMT3A-206
aaseq5491 ENST00000402667 DNMT3A-205
Q9Y6K1 ENST00000321117_MANE DNMT3A-202
g
h
SYK
NOTPRO flairiso36192
P43405-2 flairiso36191
P43405 flairiso24723
P43405 ENST00000375754_MANE SYK-204
P43405-2 ENST00000375751 SYK-203

**Supplementary Figure 9: FLAIR3 identifies tumor-biased isoform usage changes in lung cancer that impact gene function. a,** Gene expression for ERG. **b,** Isoform usage for ERG. X-axis is patient ID, above that is N (adjacent normal) and T (tumor). Center: Isoform usage (fraction of total expression of the gene). Isoforms that are different shades of the same color produce the same amino acid sequence. Grey means they are non-productive. Legend: Each isoform is identified by the UniProtKB accession (when possible), then Ensembl transcript ID and transcript name (when possible). MANE transcripts are indicated. Variants are indicated by GENIE polyPhen classification when relevant (damaging = probably_damaging), then genomic position and amino acid change (only when known to be somatic). Bottom right: Models of the isoforms. All are oriented in the 5'->3' gene direction. Thicker blocks are coding sequence (CDS). **c,** Gene expression for *ERCC1*. **d,** Isoform usage for *ERCC1*. All panels are as described in b. **e,** Gene expression for *DNMT3A*. **f,** Isoform usage for *DNMT3A*. All panels are as described in b. **g,** Gene expression for *SYK*. **h,** Isoform usage for *SYK*. All panels are as described in b.
